# A set of functionally-defined brain regions with improved representation of the subcortex and cerebellum

**DOI:** 10.1101/450452

**Authors:** Benjamin A. Seitzman, Caterina Gratton, Scott Marek, Ryan V. Raut, Nico U.F. Dosenbach, Bradley L. Schlaggar, Steven E. Petersen, Deanna J. Greene

**Author notes:** Corresponding Author: Benjamin A. Seitzman, Washington University in St. Louis, Department of Neurology 4525 Scott Ave, Box 8111, St. Louis, MO 63110, USA.

## Abstract

An important aspect of network-based analysis is robust node definition. This issue is critical for functional brain network analyses, as poor node choice can lead to spurious findings and misleading inferences about functional brain organization. Two sets of functional brain nodes from our group are well represented in the literature: (1) 264 volumetric regions of interest (ROIs) reported in Power et al., 2011 and (2) 333 cortical surface parcels reported in Gordon et al., 2016. However, subcortical and cerebellar structures are either incompletely captured or missing from these ROI sets. Therefore, properties of functional network organization involving the subcortex and cerebellum may be underappreciated thus far. Here, we apply a winner-take-all partitioning method to resting-state fMRI data to generate novel functionally-constrained ROIs in the thalamus, basal ganglia, amygdala, hippocampus, and cerebellum. We validate these ROIs in three datasets using several criteria, including agreement with existing literature and anatomical atlases. Further, we demonstrate that combining these ROIs with established cortical ROIs recapitulates and extends previously described functional network organization. This new set of ROIs is made publicly available for general use, including a full list of MNI coordinates and functional network labels.

## 1. Introduction

The brain is organized into areas that interact with one another to form distributed large-scale networks (Allman and Kaas, 1971; Felleman and Van Essen, 1991; Petersen and Sporns, 2015). Researchers studying the brain at the network level have revealed both basic principles of brain organization (Bassett and Bullmore, 2006; Honey et al., 2007; Power et al., 2011; Sporns et al., 2004; van den Heuvel and Sporns, 2011; Yeo et al., 2011) and insights into neurologic and psychiatric diseases (Corbetta and Shulman, 2011; Kim et al., 2014; Seeley et al., 2009; Sorg et al., 2007). Much of this work has borrowed concepts and tools from the field of graph theory in order to model the brain as a network (Bullmore and Sporns, 2009; Sporns, 2011). A graph is a mathematical description of a network, which comprises a set of elements (nodes) and their pairwise relationships (edges (Bondy and Murty, 1976)). Therefore, network approaches require the definition of a set of nodes, such as regions of interest (ROIs) in the case of brain networks.

Ideally, nodes should be internally coherent (e.g., functionally homogeneous) and independent, separable units (Bullmore and Bassett, 2011; Butts, 2009, 2008; Wig et al., 2011). Brain areas and their constituent components—local circuits, columns, and domains (Kaas, 2012)—display many of these properties, and thus, are suitable nodes for brain network analysis. Research efforts focused on node definition often employ data-driven techniques to parcellate the cerebral cortex into a set of ROIs meant to represent putative functionally homogeneous brain areas (Cohen et al., 2008; Craddock et al., 2012; Glasser et al., 2016; Gordon et al., 2016; Nelson et al., 2010; Power et al., 2011; Schaefer et al., 2017; Wig et al., 2013). Most such studies have used resting-state functional connectivity MRI, which measures correlations in low-frequency blood-oxygen-level-dependent (BOLD) signals across the whole brain while subjects remain awake and alert without engaging in an explicit task (Biswal et al., 1995; Gusnard and Raichle, 2001; Snyder and Raichle, 2012). While many of these existing sets of ROIs sample the cortex quite well, most approaches have under-sampled or completely omitted the subcortex and cerebellum (but see Ji et al., 2019).

The poorer representation of these structures is a limitation of previous work, as closed loop anatomical circuits connect the subcortex and cerebellum to the cortex (Woolsey et al., 2008). In addition, these structures are known to be integral for many behavioral, cognitive, and affective functions. For example, regions of the cerebellum are involved in adaptive behaviors (Thach et al., 1992), including fast adaptations, like eye-blink conditioning (Steinmetz et al., 1992; Perrett et al., 1993), as well as those that occur over longer timescales, like prism adaptation (Martin et al., 1996; Baizer et al., 1999; Morton and Bastian, 2004), and higher order cognitive functions, such as semantic processing (Fiez, 2016; Guell et al., 2018). Likewise, regions of the basal ganglia and thalamus are important for both lower level sensory and higher order cognitive functions (Alexander et al., 1986; Jones, 1985). Furthermore, subcortical structures and the cerebellum have been implicated in a variety of neurologic and psychiatric diseases. For instance, the basal ganglia are affected in several movement disorders (Greene et al., 2017, 2013; Rajput, 1993; Vonsattel et al., 1985), the hippocampus is disrupted in Alzheimer Disease (Hardy and Selkoe, 2002), the amygdala is implicated in Major Depressive Disorder (Frodl et al., 2002) and Urbach-Wiethe Disease (Siebert et al., 2003), and the cerebellum is disturbed in Schizophrenia (Andreasen et al., 1996; Bigelow et al., 2006; Brown et al., 2005; Kim et al., 2014) and Autism Spectrum Disorder (Fatemi et al., 2002), to name a few. Moreover, interactions between the cortex and both subcortical and cerebellar regions are crucial for carrying out functions in health (Bostan and Strick, 2018; Greene et al., 2014; Hwang et al., 2017; Kiritani et al., 2012) and disease (Andreasen et al., 1999; Gratton et al., 2018a; Schmahmann, 2004). Because of these interactions between multiple structures, it has been postulated that subcortical regions may have important hub-like properties for integrating brain systems (Hwang et al., 2017) and may constrain network-level topology (Bell and Shine, 2016; Garrett et al., 2018). Thus, brain network analyses should include these important regions in order to have a more complete picture of brain organization and function.

An issue potentially impeding the inclusion of these regions is that subcortical and deep cerebellar nuclei are small relative to the spatial resolution of fMRI, often occupying just a few voxels, whereas brain areas in the cerebral cortex (e.g. Area V1) are typically larger. Furthermore, depending on the acquisition sequence, these regions may have lower signal quality (Ojemann et al., 1997) or, especially for the cerebellum, may be captured incompletely. Finally, most existing techniques for parcellating the brain into areas, such as gradient-based techniques (Cohen et al., 2008; Gordon et al., 2016; Nelson et al., 2010; Wig et al., 2013), were designed for the cortical surface, making them less easily applied to structures where surface-based mapping is less appropriate (basal ganglia, thalamus), prone to error (medial temporal lobe) (Wisse et al., 2014), or less well-established (cerebellum). Despite these difficulties, inclusion of the subcortex and cerebellum is crucial to properly represent the brain as a network. While there are existing anatomical atlases of the subcortex (Morel, 2013) and cerebellum (Diedrichsen et al., 2009), functionally defined regions may complement anatomical ones and provide a better correspondence to functionally defined cortical areas and task-based measures from fMRI.

Our lab previously published two (now widely used) sets of ROIs: (1) 264 volumetric ROIs (Power et al., 2011) and (2) 333 surface-based cortical parcels (Gordon et al., 2016). The first was created via combined task fMRI meta-analysis and resting-state functional correlation mapping, and the second was created via a gradient-based parcellation of resting-state fMRI data. These two ROI sets sample the cortex well, representing a diverse set of brain areas that can be organized into functional networks. Many investigators have used them to describe functional brain organization in a variety of healthy samples (Power et al., 2013; Zanto and Gazzaley, 2013), lifespan cohorts (Baniqued et al., 2018; Gallen et al., 2016; Gu et al., 2015; Nielsen et al., 2018; Rudolph et al., 2017), as well as populations with neurologic and psychiatric diseases (Gratton et al., 2018a; Greene et al., 2016; Sheffield et al., 2015; Siegel et al., 2018). However, the first set (264 volumetric ROIs) under-samples subcortical and cerebellar structures, as only 17 ROIs are non-cortical, and the second set (333 parcels) is restricted to the cortex only, similar to other popular ROI sets, e.g. (Glasser et al., 2016; Yeo et al., 2011).

The goal of the current study was to expand these ROI sets to better represent subcortical and cerebellar structures. Novel ROIs were created in the thalamus, basal ganglia, and cerebellum by use of a data-driven, winner-take-all partitioning technique that operates on resting-state fMRI data (Choi et al., 2012; Greene et al., 2014; Zhang et al., 2010). Additional ROIs were generated in the amygdala and hippocampus, and all ROIs were validated via several criteria. Finally, we characterized whole-brain functional network organization using these refined subcortical and cerebellar ROIs combined with previously established cortical ROIs. The fully updated set of ROIs is made publicly available for general use, including a list of coordinates and consensus functional network labels, at https://greenelab.wustl.edu/data_software.

## 2. Material and Methods

### 2.1. Primary dataset-WashU 120

#### 2.1.1. Dataset characteristics

The primary dataset used in this study has been described previously (Power et al., 2011). Eyes-open resting-state fMRI data were acquired from 120 healthy, right-handed, native English speaking, young adults (60 F, age range 18-32, mean age 24.7). Subjects were recruited from the Washington University community and screened with a self-report questionnaire. Exclusion criteria included no current or previous history of neurologic or psychiatric diagnosis as well as no head injuries resulting in a loss of consciousness for more than 5 minutes. Informed consent was obtained from all participants, and the Washington University Internal Review Board approved the study. The data are available at https://legacy.openfmri.org/dataset/ds000243/.

#### 2.1.2. Data acquisition

A Siemens MAGNETOM Tim TRIO 3.0T MRI scanner and a 12 channel Head Matrix Coil were used to obtain T1-weighted (MP-RAGE, 2.4s TR, 1×1×1mm voxels) and BOLD contrast sensitive (gradient echo EPI, 2.5s TR, 4×4×4mm voxels) images from each subject. The mean amount of BOLD data acquired per subject was 14 minutes (336 frames, range = 184-729 frames). Subjects were instructed to fixate on a black crosshair presented at the center of a white background. See Power et al., 2011 for full acquisition details.

#### 2.1.3. Preprocessing

The first 12 frames (30 seconds) of each functional run were discarded to account for magnetization equilibrium and an auditory evoked response at the start of the EPI sequence (Laumann et al., 2015). Slice timing correction was applied first. Then, the functional data were aligned to the first frame of the first run using rigid body transforms, motion corrected (3D-cross realigned), and whole-brain mode 1000 normalized (Miezin et al., 2000). Next, the data were resampled (3 cubic mm voxels) and registered to the T1-weighted image and then to a WashU Talairach atlas (Ojemann et al., 1997) using affine transforms in a one-step operation (Smith et al., 2004).

Additional preprocessing of the resting-state BOLD data was applied to remove artifacts (Ciric et al., 2017; Power et al., 2014). Frame-wise displacement (FD) was calculated as in Power et al., 2012, and frames with FD greater than 0.2 mm were censored. Uncensored segments with fewer than 5 contiguous frames were censored as well (mean +/- std frames retained = 279 +/- 107). All censored frames were interpolated over using least squares spectral estimation (Hocke and Kämpfer, 2009; Power et al., 2014). Next, the data were bandpass filtered from 0.009-0.08 Hz and nuisance regression was implemented. The regression included 36 regressors: the whole-brain mean, individually defined white matter and ventricular CSF signals, the temporal derivatives of each of these regressors, and an additional 24 movement regressors derived by expansion (Friston et al., 1996; Satterthwaite et al., 2012; Yan et al., 2013). FreeSurfer 5.3 automatic segmentation was applied to the T1-weighted images to create masks of the gray matter, white matter, and ventricles for the individual-specific regressors (Fischl et al., 2002). Finally, the data were smoothed with a Gaussian smoothing kernel (FWHM = 6 mm, sigma = 2.55).

At the end of all processing, each censored/interpolated frame was removed from the time series for all further analyses.

### 2.2. Secondary dataset-HCP 80

#### 2.2.1. Dataset characteristics

Due to a partial cutoff of cerebellar data in over half of the subjects in the primary dataset (outside of the field of view), an independent secondary dataset was used to supplement analyses related to the cerebellum. Since the cerebellum was not cutoff in every subject in the primary dataset, we were able to create a cerebellar portion of the group average matrix derived from just those subjects with full cerebellar coverage. We used data from 80 unrelated individuals from the Human Connectome Project (HCP) 500 Subject Release (40F, age range 22-35, mean age 28.4) who had high-quality (low-motion) data, described previously (Gordon et al., 2017a). All HCP data are available at https://db.humanconnectome.org.

#### 2.2.2. Data acquisition

A custom Siemens SKYRA 3.0T MRI scanner and a custom 32 channel Head Matrix Coil were used to obtain high-resolution T1-weighted (MP-RAGE, 2.4s TR, 0.7×0.7×0.7mm voxels) and BOLD contrast sensitive (gradient echo EPI, multiband factor 8, 0.72s TR, 2×2×2mm voxels) images from each subject. The HCP used sequences with left-to-right and right-to-left phase encoding, with a single RL and LR run on each day for two consecutive days for a total of four runs (Van Essen et al., 2012). Thus, for symmetry, the BOLD time series from each subject’s best (most frames retained after censoring) LR run and their best RL run were concatenated together.

#### 2.2.3. Preprocessing

The preprocessing steps were the same as those detailed in Section 2.1.3 except for the following: (1) the first 41 frames (29.52 seconds) of each run were discarded, (2) no slice timing correction was applied, (3) field inhomogeneity distortion correction was applied (using the mean field map), (4) the data were not resampled (they were collected at 2 cubic mm isotropic voxels), and (5) the Gaussian smoothing kernel was smaller (FWHM = 4 mm, sigma = 1.7). The first two changes are due to the increased temporal resolution of the HCP data acquisition (0.72s TR) and the last two changes are due to the increased spatial resolution of HCP data acquisition (Glasser et al., 2013). Distortion correction was not applied to the primary dataset because field maps were not collected in most participants. In addition, the increased temporal resolution caused respiration artifacts to alias into the FD trace (Fair et al., 2018; Siegel et al., 2017). Thus, FD values were filtered with a lowpass filter at 0.1 Hz and the filtered FD threshold was set at 0.1 mm (mean +/- std frames retained = 2236 +/- 76).

For the purpose of the winner-take-all partitioning of the secondary dataset (described in section 2.4), a CIFTI was created for each subject. Thus, preprocessed cortical BOLD time series data (from the secondary dataset only) were mapped to the surface, following the procedure of Gordon et al., 2016, and combined with volumetric subcortical and cerebellar data in the CIFTI format (Glasser et al., 2013; Gordon et al., 2016).

At the end of all processing, each censored/interpolated frame was removed from the time series for all further analyses.

### 2.3. Validation dataset-MSC

#### 2.3.1. Dataset characteristics

Since the primary and secondary datasets were used to create the subcortical and cerebellar ROIs (described in section 2.5), results for functional network community assignment (described in section 2.7) were validated with a third independent dataset, the Midnight Scan Club (MSC), described previously (Gordon et al., 2017b). These data are available at https://openneuro.org/datasets/ds000224/versions/00002. The MSC dataset consists of 5 hours of resting-state BOLD data from each of 10 individuals (5 F, age range 24-34, mean age 29) over a two-week period.

#### 2.3.2. Data acquisition

The same scanner, head coil, and acquisition parameters described in Section 2.1.2 were used to for the MSC. However, a single resting-state run lasting 30 minutes was collected on 10 separate days. Each scan was acquired starting at midnight (Gordon et al., 2017b).

#### 2.3.3. Preprocessing

For each subject, all runs were concatenated together in the order that they were collected. The initial preprocessing steps were the same as those detailed in Section 2.1.3 except for the following: (1) the functional images were registered to the average T2-weighted anatomical image (4 were collected per subject), then to the average T1-weighted anatomical image (4 were collected per subject), and finally to the Talairach atlas, (2) field inhomogeneity distortion correction was applied (using the mean field map), and (3) one subject (MSC08) was excluded due to a substantial amount of low-quality data and self-reported sleeping during acquisition, as detailed previously (Gordon et al., 2017b; Laumann et al., 2016).

Additional preprocessing followed Raut and colleagues (Raut et al., 2019). Again, FD was used to exclude high-motion frames; however, due to respiratory artifacts affecting the realignment parameters (Power et al., 2018; Siegel et al., 2017), a lowpass filter (0.1 Hz) was applied to those parameters before calculation of FD. Consequently, the threshold for frame censoring was lowered to 0.1mm. Frames with outstanding (>2.5 standard deviations above the mode computed across all runs) DVARS values (as calculated in Power et al., 2012) were also excluded. All censored frames were linearly interpolated, and then bandpass filter (0.005-0.1 Hz) was applied.

Finally, component-based nuisance regression was implemented. Individual-specific FreeSurfer 6.0 segmentation was used to define masks of the gray matter, white matter, and ventricles. A mask of extra-axial (or edge (Patriat et al., 2015)) voxels was also created by thresholding the temporal standard deviation image (>2.5%) that excluded the eyes and a dilated whole-brain mask. BOLD data was extracted from all voxels in each mask (separately), and dimensionality reduction was applied as in CompCor (Behzadi et al., 2007). The number of components retained was determined independently for each mask such that the condition number (i.e., the maximum eigenvalue divided by the minimum eigenvalue) was greater than 30. All retained components were submitted to a regressors matrix that also included the 6 realignment parameters. To avoid collinearity, singular value decomposition was applied to the regressors covariance matrix. Components of this decomposition were retained up to an upper limit (condition number >=250). Then, all of the final retained components, the whole-brain mean, and its temporal derivative were regressed from the BOLD time series (Raut et al., 2019).

At the end of all processing, each censored/interpolated frame was removed from the time series for all further analyses.

### 2.4. Winner-take-all partitioning of the subcortex and cerebellum

In order to identify functional subdivisions within subcortical structures and the cerebellum, a winner-take-all partitioning technique was applied to the basal ganglia, thalamus, and cerebellum, as previously described (Greene et al., 2014). Past applications of this winner-take-all approach have yielded results consistent with known connectivity from the animal literature (Buckner et al., 2011; Choi et al., 2012; Fair et al., 2010; Greene et al., 2014; Zhang et al., 2008).

Briefly, the mean resting-state time series were extracted from each of 11 previously defined cortical networks (Power et al., 2011): default mode, frontoparietal, cinguloopercular, salience, dorsal attention, ventral attention, visual, auditory, somatomotor dorsal, somatomotor lateral, and orbitofrontal. This subset of networks (from the original 15 described in Power et al., 2011) was selected on the basis of being previously well characterized and validated by multiple methods (see Yeo et al., 2011; Greene et al., 2014). In order to remove the shared variance among cortical networks thereby increasing specificity of the subcortico-cortical and cerebello-cortical correlations, partial correlations were then calculated between the time series from each cortical network and the resting-state time series from each subcortical or cerebellar gray matter voxel (e.g., for each cortical network and subcortical voxel, a residual correlation was computed after partialling out the signal from the other cortical networks). Each voxel was then assigned to the network with which it correlated most in a winner-take-all fashion (Buckner et al., 2011; Choi et al., 2012; Greene et al., 2014; Zhang et al., 2010), generating a functional partition of subcortical and cerebellar structures.

### 2.5. ROI creation

Spherical ROIs (diameter = 8mm) were placed in the (volumetric) center of each of the winner-take-all partitions in the basal ganglia, thalamus, and cerebellum. Then, the ROIs were manually adjusted such that (1) all ROIs included only gray matter voxels and (2) no ROIs had any overlapping voxels. If an ROI did not fit entirely within a single winner-take-all partition, it was excluded. Two additional ROIs (one per hemisphere) were added to the center of the amygdala, since the entire structure was assigned to a single network (default mode) via the winner-take-all approach. The winner-take-all approach also assigned the entire hippocampus to a single network (default mode).

However, given previous evidence for distinct functional connectivity profiles for the anterior and posterior portions of the hippocampus (Kahn et al., 2008), we added four ROIs (two per hemisphere) to sample the anterior and posterior hippocampus. In total, 34 subcortical and 27 cerebellar ROIs were created.

These new subcortical and cerebellar ROIs were then combined with two previously described sets of cortical ROIs from our lab, as follows:

#### ROI Set 1 (Power264 + new)

Spherical cortical ROIs were used from the 264 volumetric ROIs reported in (Power et al., 2011). Four of these ROIs in the medial temporal lobe (two per hemisphere) were removed (Talairach coordinates: (−20, −24, −18), (17, −30, −15), (−25, −41, −8), (26, −39, −11)) and replaced by the four new hippocampus ROIs, due to some overlapping voxels. In addition, the 17 subcortical and cerebellar ROIs from the original 264 were replaced by 55 new subcortical and cerebellar ROIs. Finally, the 2 new amygdala ROIs were added. Thus, ROI Set 1 is composed of 239 cortical, 34 subcortical (including the amygdala and hippocampus), and 27 cerebellar volumetric ROIs, for a total of 300 ROIs.

#### ROI Set 2 (Gordon333 + new)

ROI set 2 was generated by combining the 333 surface-based cortical parcels (Gordon et al., 2016) with the newly generated subcortical and cerebellar ROIs. Thus, ROI Set 2 is composed of 333 surface-based cortical parcels and 34 subcortical (including the amygdala and hippocampus) and 27 cerebellar volumetric ROIs, for a total of 394 ROIs. For all analyses using this ROI set, we utilized the center of each cortical parcel projected into volumetric atlas space (Gordon et al., 2016). The parcels in this format are publicly available at https://sites.wustl.edu/petersenschlaggarlab/parcels-19cwpgu/.

### 2.6. Seedmaps and consensus functional network communities for each ROI

#### 2.6.1. Seedmaps

To validate the winner-take-all assignments of voxels used for ROI placement, we first conducted seedmap analyses to examine how each ROI was correlated with every other gray matter voxel. A seedmap represents the pattern of correlations between the mean BOLD time series from a given ROI and all other gray matter voxels in the brain. We generated group-average seedmaps for both ROI Sets and each dataset (primary, secondary, validation). The preprocessed BOLD time series for each gray matter voxel within each ROI were averaged together (after removing censored and interpolated frames). Then, the Pearson correlation between each new ROI and every other gray matter voxel in the brain was computed for each subject. The subject-specific maps were Fisher transformed, averaged together, and inverse Fisher transformed.

#### 2.6.2. Correlation matrices

We generated correlation matrices to examine the community structure of the new ROIs. A correlation matrix is the set of all possible pairwise correlations between mean BOLD time series from each ROI organized into a symmetric matrix (since correlations are undirected). We computed correlation matrices for both ROI Sets and each dataset (primary, secondary, validation). The preprocessed BOLD time series for each gray matter voxel within each ROI were averaged together (after removing censored and interpolated frames). Then, the Pearson correlation between every pair of ROIs was computed to create a 300 x 300 (ROI Set 1) and 394 x 394 (ROI set 2) correlation matrix for each subject. Matrices were individually Fisher Z transformed, all matrices were averaged together (within each ROI set and dataset; thus, six group-average matrices were created in total-one 300 x 300 and one 394 x 394 for each of the WashU 120, HCP 80, and MSC 9), and finally, inverse Fisher transformed.

#### 2.6.3. Community detection

To determine the functional network membership of each ROI, an information-theoretic community detection algorithm was implemented (InfoMap (Rosvall and Bergstrom, 2008)). InfoMap requires a sparse matrix, so an edge density threshold was applied to the correlation matrices. The networks (correlation matrices) were thresholded until only the strongest X percent of edges remained. All retained edges maintained their correlation value or weight (i.e., the networks were not binarized). We ran InfoMap over a range of thresholds (X = 2-10% inclusive, with a 1% step increment, following Power et al. (2011)).

In general, the magnitude of BOLD correlations between the cortex and the subcortex, the cortex and the cerebellum, and the subcortex and the cerebellum is substantially weaker than within-structure (and particularly, cortico-cortical) correlations. The primary reasons for this are likely distance from the head matrix coil and signal dropout due to sinuses. For instance, in the primary dataset, off-diagonal (between-structure) correlations from the subcortex and cerebellum account for 40% of the weakest decile of correlations (i.e., the 10% of correlations closest to 0), even though the subcortex and cerebellum account for only 23% of all ROIs. Therefore, in order to ensure that between-structure correlations were included, structure-specific thresholding was used (Marek et al., 2018). The correlation matrix was separated into cortical, subcortical, and cerebellar components (e.g., the subcortical component is every entry in each row corresponding to any subcortical ROI) and the edge density thresholds were applied to each component separately. Thus, if a 2% structure-specific edge density was applied to the matrix, the top 2% of cortical, top 2% of subcortical, and top 2% of cerebellar correlations (excluding diagonal entries) were extracted and all other correlations were set to 0.

#### 2.6.4. Consensus network procedure

Consensus functional network communities were determined in a semi-automated, multistep process. First, a weighting procedure was applied across InfoMap thresholds. For the 2% and 3% thresholds the weight was 5, for the 4% and 5% thresholds the weight was 3, and the weight was 1 for all other thresholds. These weights were chosen to bias the consensus solution to have approximately 17 networks on the basis of work from Yeo and colleagues (Yeo et al., 2011). Since smaller networks tend to be observed at sparser thresholds, those thresholds contribute more weight than the denser thresholds. For each ROI (independently), the InfoMap-determined community at each threshold was noted, taking the weights into account, and the highest weighted community was assigned as the consensus.

After this automated consensus procedure, authors BAS, CG, and DJG reviewed the community assignment of each new subcortical and cerebellar ROI. In ambiguous cases (e.g., an even split in assignment across thresholds), we consulted literature describing the anatomy and function of that brain region. In cases when the InfoMap and the winner-take-all assignments differed, we adjudicated between the two using a previously described template-matching algorithm (from (Gordon et al., 2017a)). Briefly, a seedmap was generated for each ROI, and binarized such that all correlation values greater than 0 were set to 1 (all others set to 0). Then, the cortical portion of this binary seedmap was compared to 14 network templates (binary representations of whole-network seedmaps defined *a priori*) via the dice coefficient. Each ROI received the network label for which the dice overlap was maximal. There were 24 ROIs that required adjudication in this way.

All cortical ROIs retained their original assignment from published works (Power et al., 2011 for ROI Set 1 and from Gordon et al., 2016 for ROI Set 2) unless there was strong evidence to overturn the original. Specifically, if an ROI in the present InfoMap solution received the same assignment across all thresholds and that assignment was distinct from the original, then the ROI was assigned to the novel network community (detailed in Section 3.3). Furthermore, 5 ROIs originally assigned to the salience network were reassigned to the cingulo-opercular network. We made this change because (1) the ROIs showed profiles intermediate between salience and cinguloopercular assignments and (2) previously published studies revealed that these brain regions demonstrate task-evoked activity consistent with the cingulo-opercular network (Dosenbach et al., 2006; Dubis et al., 2016; Gratton et al., 2018b, 2017; Neta et al., 2014).

#### 2.6.5. Validation of ROIs and consensus networks

The primary and secondary datasets were used to create the subcortical and cerebellar ROIs, respectively. The validation dataset (MSC) was used to test the validity of the consensus functional network communities in both cases. The network community assignment for each ROI was compared across all datasets, and discrepancies were noted. Further, consensus networks were compared with those from previously published literature including the Morel anatomical atlas of the subcortex and the SUIT anatomical atlas of the cerebellum. Additionally, the winner-take-all assignments were compared between split-halves of the primary dataset. Finally, we measured the degree of confidence in the “winning” network for each subcortical and cerebellar voxel by calculating the difference in functional connectivity between the winning and second place network assignments, as in Marek et al., 2018. This analysis was conducted with the primary dataset (WashU 120) for the basal ganglia and thalamus, and with the validation dataset (MSC) for the cerebellum. We examined the location of each ROI with respect to this estimation of confidence in the winner-take-all assignments (SI Figure 1).

### 2.7. Accounting for ROIs with strong functional connectivity to multiple networks

The InfoMap and winner-take-all approaches each yield a single network solution for each ROI. However, there is evidence that regions within the subcortex connect with multiple functional networks, as previous studies have identified zones of integration within the basal ganglia and thalamus (Greene et al., under review; Garrett et al., 2018; Hwang et al., 2017). Such integration may be a source of low-confidence winner-take-all assignments (SI Figure 1). To address this issue, we used the validation dataset (MSC) to identify ROIs that overlap with “integrative” voxels. As in Greene et al., (under review), a voxel was considered integrative if its correlation with any network was within 66.7% of its correlation with the “winning” network. Note that identification of integrative voxels is highly similar across thresholds (within 50-75% of the “winning” network) and with different methodologies (e.g., a method based on effect size). Then, we calculated the percent of voxels within each non-cortical ROI that overlapped with integrative voxels. ROIs with a majority of overlapping voxels (>50%) were flagged as “integrative.”

### 2.8. Spring-embedded graphs and participation coefficient

To visualize the community structure of networks in an abstract graph space, spring-embedded graphs were created. The networks (correlation matrices) were thresholded in the same way as in Section 2.6.3, and the resulting matrices were submitted to a physical model of connected springs (the Kamada-Kawai algorithm, as used in Power et al., 2011). Correlations between pairs of ROIs were modeled as force constants between connected springs such that strongly correlated ROIs were “pulled” close to one another. ROIs were colored according to their consensus functional network community or their anatomical location.

To quantify the degree to which an ROI plays a hub-like role in the network, the participation coefficient of each ROI was computed across (structure-specific) edge density thresholds between 5 and 25%. Participation coefficient was calculated as defined for weighted networks in Rubinov and Sporns, 2010 using code from the Brain Connectivity Toolbox (Rubinov and Sporns, 2010).

## 3. Results

### 3.1. Subcortical and cerebellar ROIs

The final set of subcortical and cerebellar ROIs overlaid onto the winner-take-all partitions are displayed in Figures 1 and 2, respectively. The winner-take-all partitions were similar to previously published partitions for the basal ganglia (Choi et al., 2012; Greene et al., 2014), thalamus (Hwang et al., 2017), and cerebellum (Buckner et al., 2011), and showed good split-half replication (dice overlap of 61.5% in the thalamus and 60.1% in the basal ganglia; SI Figure 2). Many ROIs outside the cortex agree with anatomical divisions from previously established subcortical (Morel, 2013) and cerebellar (Diedrichsen et al., 2009) atlases, as shown in Figure 3. ROIs that do not show perfect correspondence with anatomical parcels may reflect discrepancies between anatomical and functional division of these structures, potentially due to finer parcellations in the anatomical atlases. A majority of the ROIs were contained within high confidence winner-take-all parcels, as assessed in Marek et al., 2018 (SI Figure 1).

**Figure 1:**
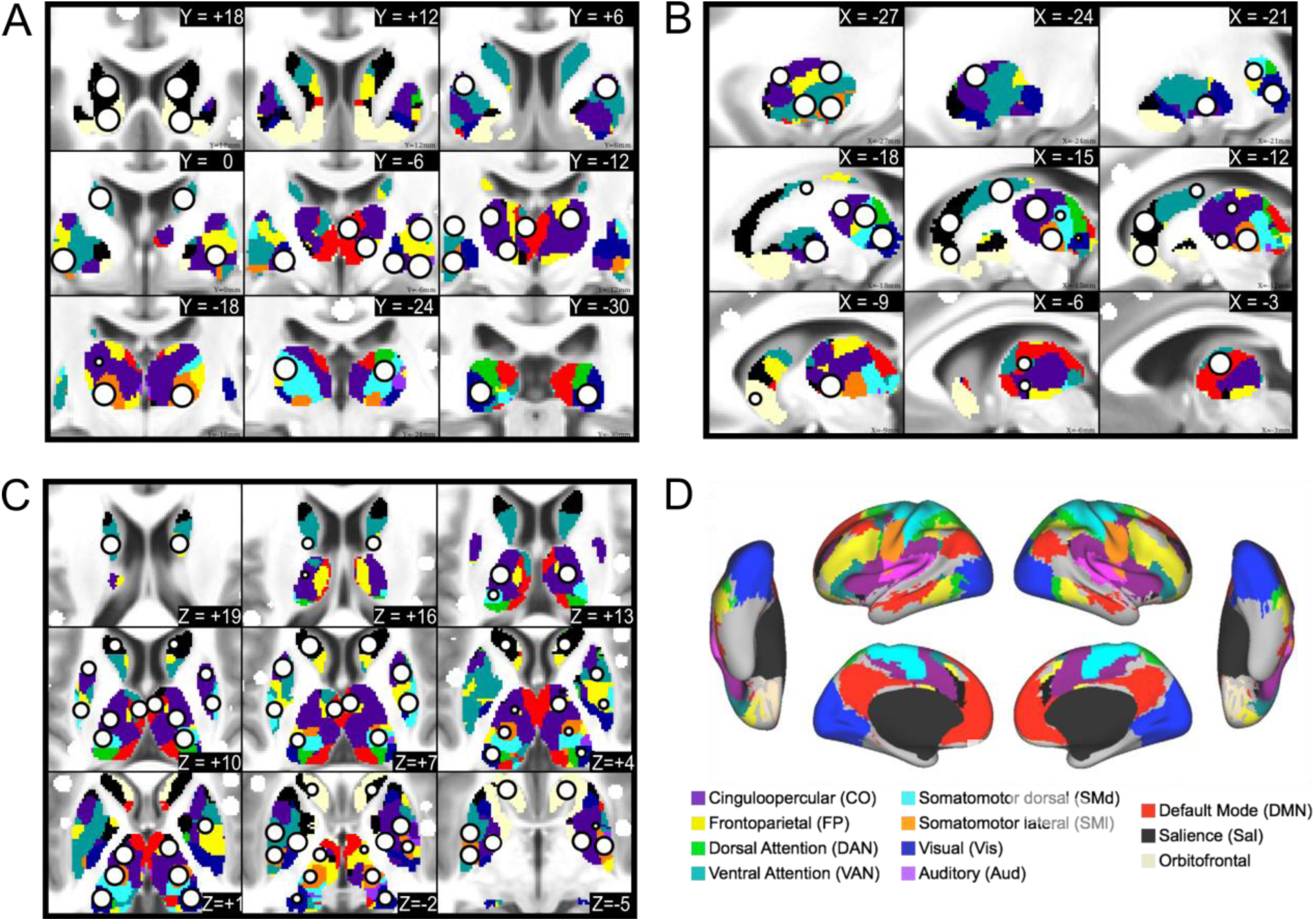
Subcortical ROIs. The new ROIs (white circle with black outline) are displayed in serial coronal (A), sagittal (B), and axial (C) sections of the thalamus and basal ganglia, with the cortical functional networks for reference (D). The ROIs are overlaid on top of the voxel-wise winner-take-all partitions.

**Figure 2:**
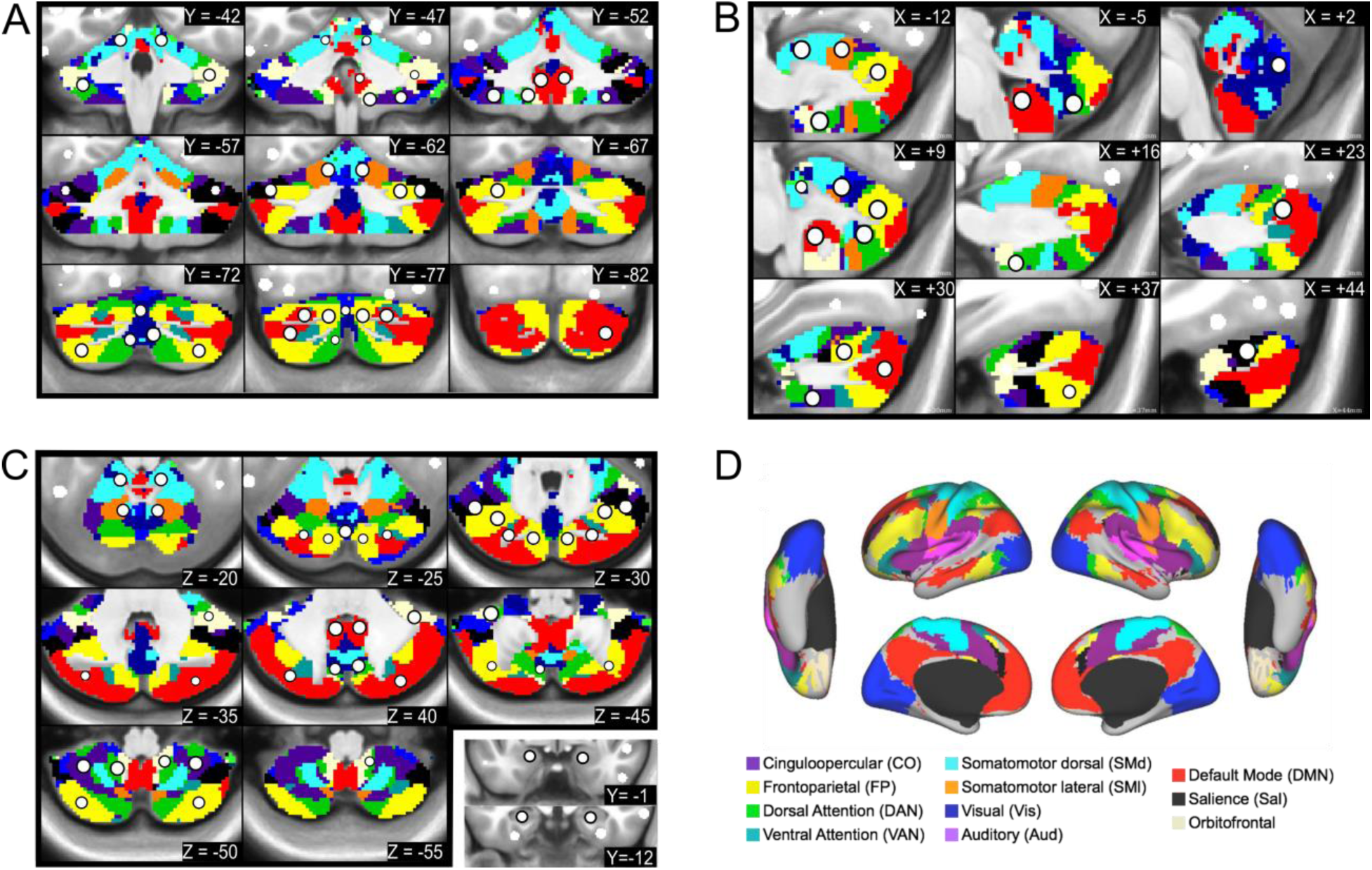
Cerebellar ROIs. The new ROIs (white circle with black outline) are displayed in serial coronal (A), sagittal (B), and axial (C) sections of the cerebellum, with the cortical functional networks for reference (D). The ROIs are overlaid on top of the voxel-wise winner-take-all partitions. ROIs in the amygdala and anterior hippocampus are overlaid on anatomical coronal sections in the bottom right panel of C.

**Figure 3:**
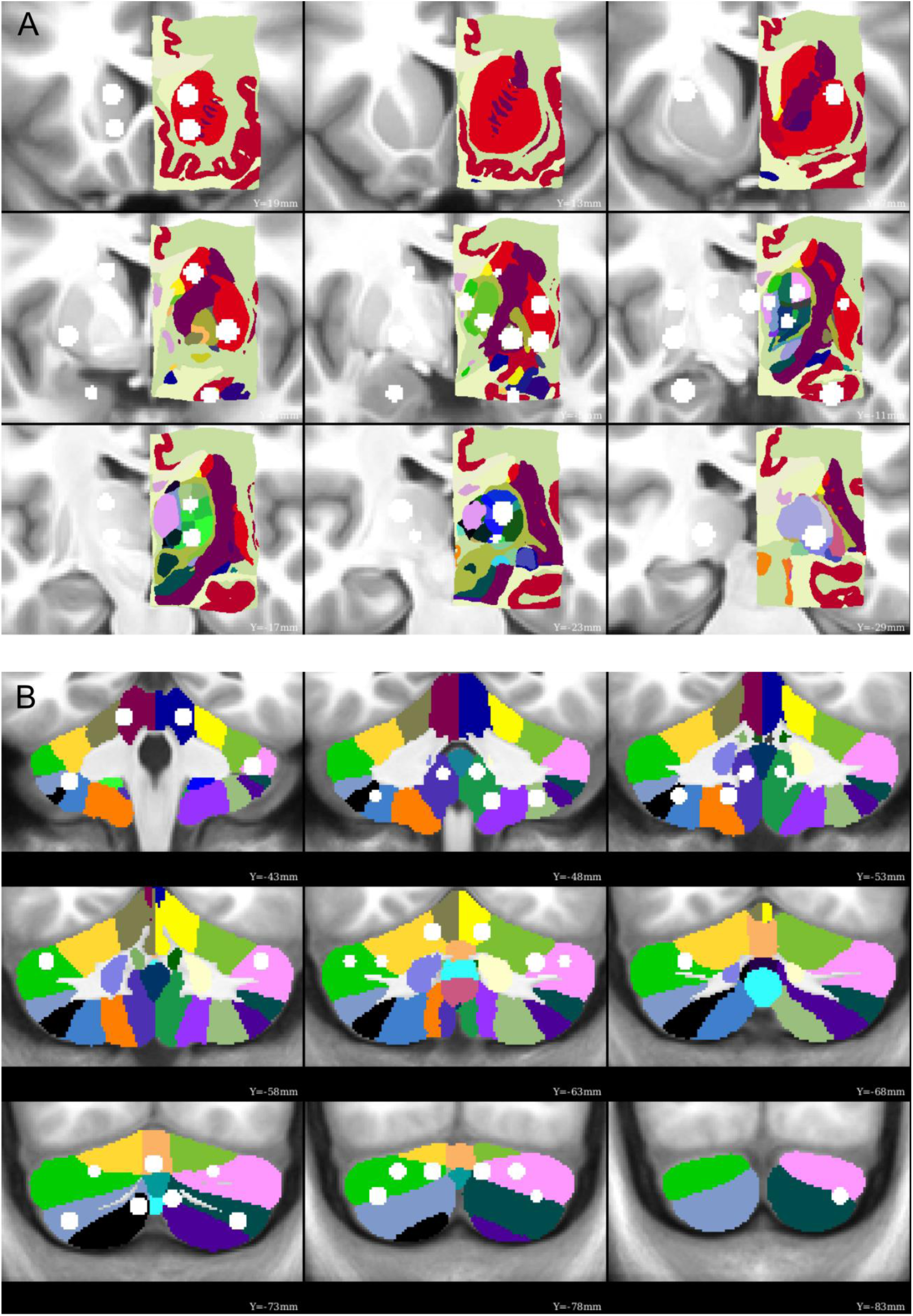
Functionally-defined ROIs overlaid onto anatomical parcellations. Many of the subcortical and cerebellar ROIs are contained within a single anatomical parcel from the Morel atlas of the subcortex (A) and the SUIT atlas of the cerebellum (B), indicating good agreement between the current functional parcellation. A few ROIs overlap multiple anatomical parcels (e.g., dorsolateral thalamus, right posterior cerebellum), which may be a consequence of a finer parcellation than is possible with the current fMRI data.

The 34 subcortical ROIs sampled the following anatomical structures (bilaterally): the head and tail of the caudate; anterior dorsal, posterior dorsal, anterior ventral, and posterior ventral putamen; the globus pallidus (internus and externus combined); the ventral striatum (i.e., nucleus accumbens); the amygdala (nuclei not distinguished); anterior and posterior hippocampus; and regions in the thalamus. The locations of the thalamic ROIs included the following nuclei and surrounding territory (the resolution of our data was not fine enough to delineate precise thalamic nuclei): medio-dorsal (MD), latero-dorsal (LD), ventro-anterior (VA), ventro-lateral (VL), ventro-postero-lateral (VPL), and lateral geniculate nucleus (LGN)-pulvinar. The 27 cerebellar ROIs sampled the vestibulo-, spino-, and cerebro-cerebellum, including the cerebellar vermis, classical motor cerebellar cortex, and cerebellar association cortex (Woolsey et al., 2008).

### 3.2. Correlation structure replicates across datasets

Exemplar seedmaps from the new ROIs for the primary dataset are displayed in Figure 4 and the group-average correlation matrices for all datasets using ROI Set 1 are displayed in Figure 5. The correlation matrices using ROI Set 2 are displayed in SI Figure 3. The seedmaps were comparable to previously published maps (Figure 4). The matrices were quite similar across datasets (r_120,HCP_ = 0.90, r_120,MSC_ = 0.93, r_HCP,MSC_ = 0.87), with results from the primary dataset replicating best in the validation (MSC) dataset. However, in the secondary (HCP) dataset, there was approximately 0 correlation between subcortical ROIs and all other ROIs, including homotopic subcortical ROI pairs. The likely reason for this difference is due to poor temporal signal-to-noise ratio in the subcortex of HCP data (Ji et al., 2019), which we demonstrate here in SI Figure 4. Thus, we excluded the secondary dataset from all further analyses.

**Figure 4:**
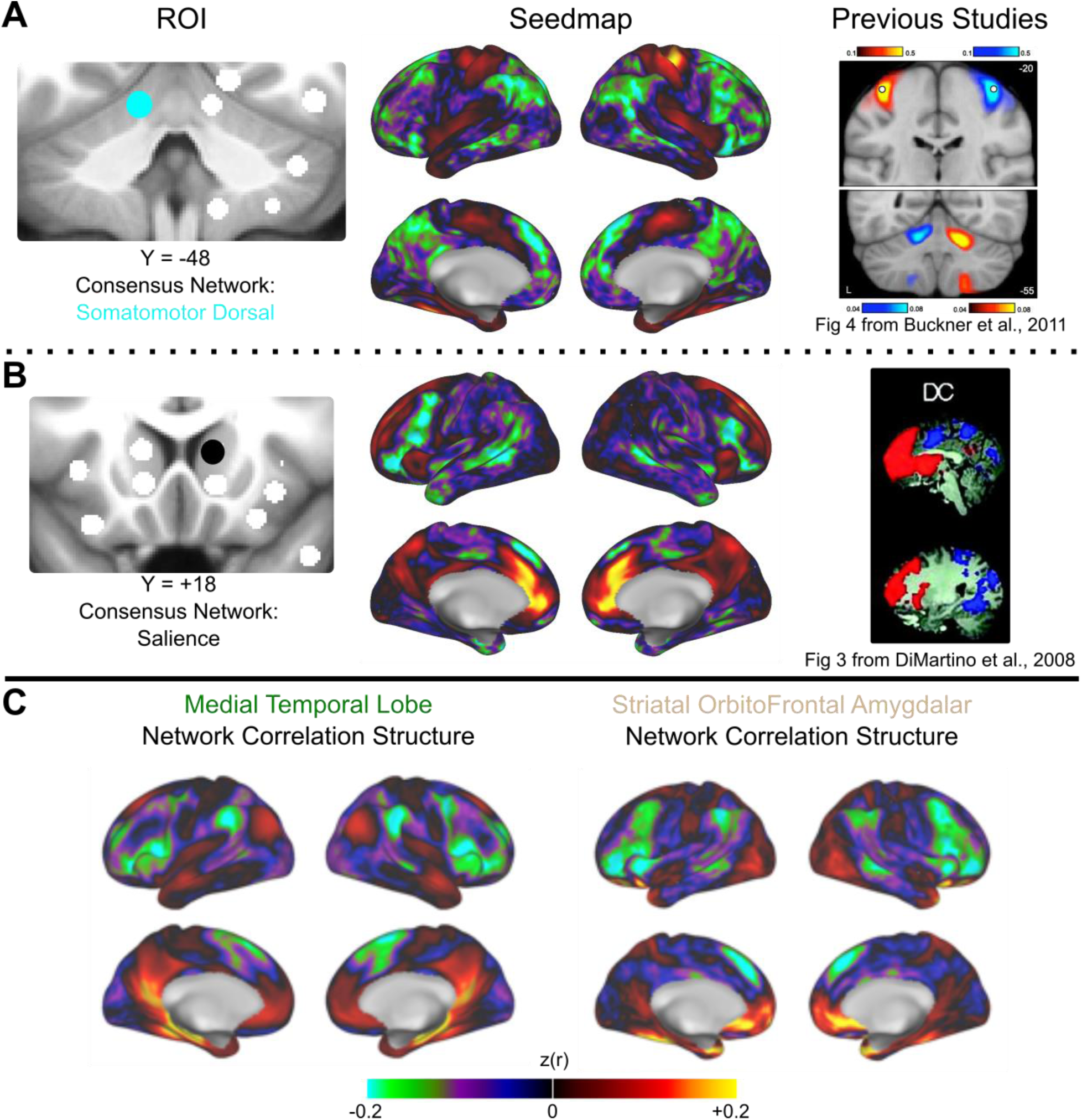
Exemplar seedmaps for the new ROIs. Functional correlation seedmaps are shown for an exemplar ROI in the cerebellum (A) and dorsal striatum (B). The consensus functional network assignment of each ROI is represented by its color (left column). Seedmaps display the correlations between the mean BOLD signal from the ROI in question and the BOLD signal from every other gray matter voxel (middle column). Results were similar to comparable seedmaps from previously published studies (right column). Images from Buckner, R.L. et al., 2011. The organization of the human cerebellum estimated by intrinsic functional connectivity. *Journal of Neurophysiology* 106 (5) 2322–2345; and, Di Martino, A. et al., 2008. Functional Connectivity of Human Striatum: A Resting State fMRI Study. *Cerebral Cortex* 18 (12), 2735–2747 reproduced with permission from The American Psychological Society and Oxford University Press. (C) Correlation structure for the Medial Temporal Lobe (left) and Striatal OrbitoFrontal Amygdalar (right) networks are displayed (see Section 3.3).

**Figure 5:**
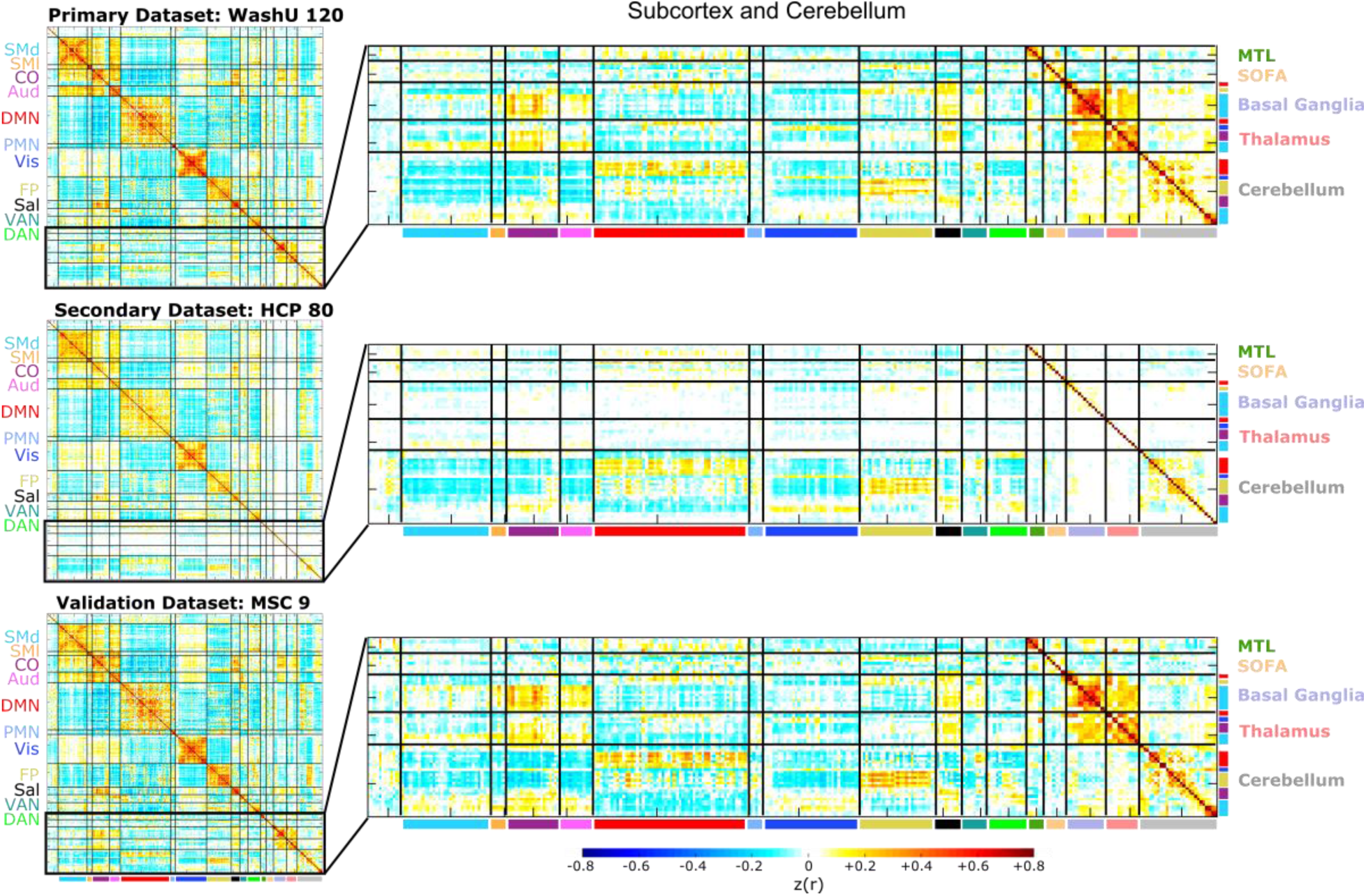
Correlation matrices are similar across datasets. The full (300 x 300) correlation matrices for ROI Set 1 are displayed for each dataset in the left column, and zoomed-in versions of the subcortical and cerebellar portions of the matrices are displayed in the right column (the corresponding images for ROI Set 2 are shown in SI Figure 3). The cortical portion of the correlation matrix is sorted by functional network community, whereas the subcortical and cerebellar portions are sorted first by anatomical structure (i.e., basal ganglia, thalamus, and cerebellum) and then by functional network community (within each structure). The matrices are similar to one another (e.g., the correlation between the primary and validation datasets is 0.93), except for the subcortical portion of the secondary dataset (HCP-Human Connectome Project). We observed poor temporal signal-to-noise in subcortical HCP data (SI Figure 4). The first row and column of the matrices correspond to unlabeled regions (i.e., InfoMap was unable to assign these ROIs to a network, similar to Power et al., 2011 and Gordon et al., 2016).

### 3.3. Functional network organization using the expanded ROI Set

We used a data-driven community detection algorithm (InfoMap) on weighted networks to determine the functional network community membership of the expanded set of ROIs (Rosvall and Bergstrom, 2008). The results of this analysis are displayed in Figure 6. Communities are shown for all tested edge density thresholds alongside the consensus network communities (see section 2.6).

**Figure 6:**
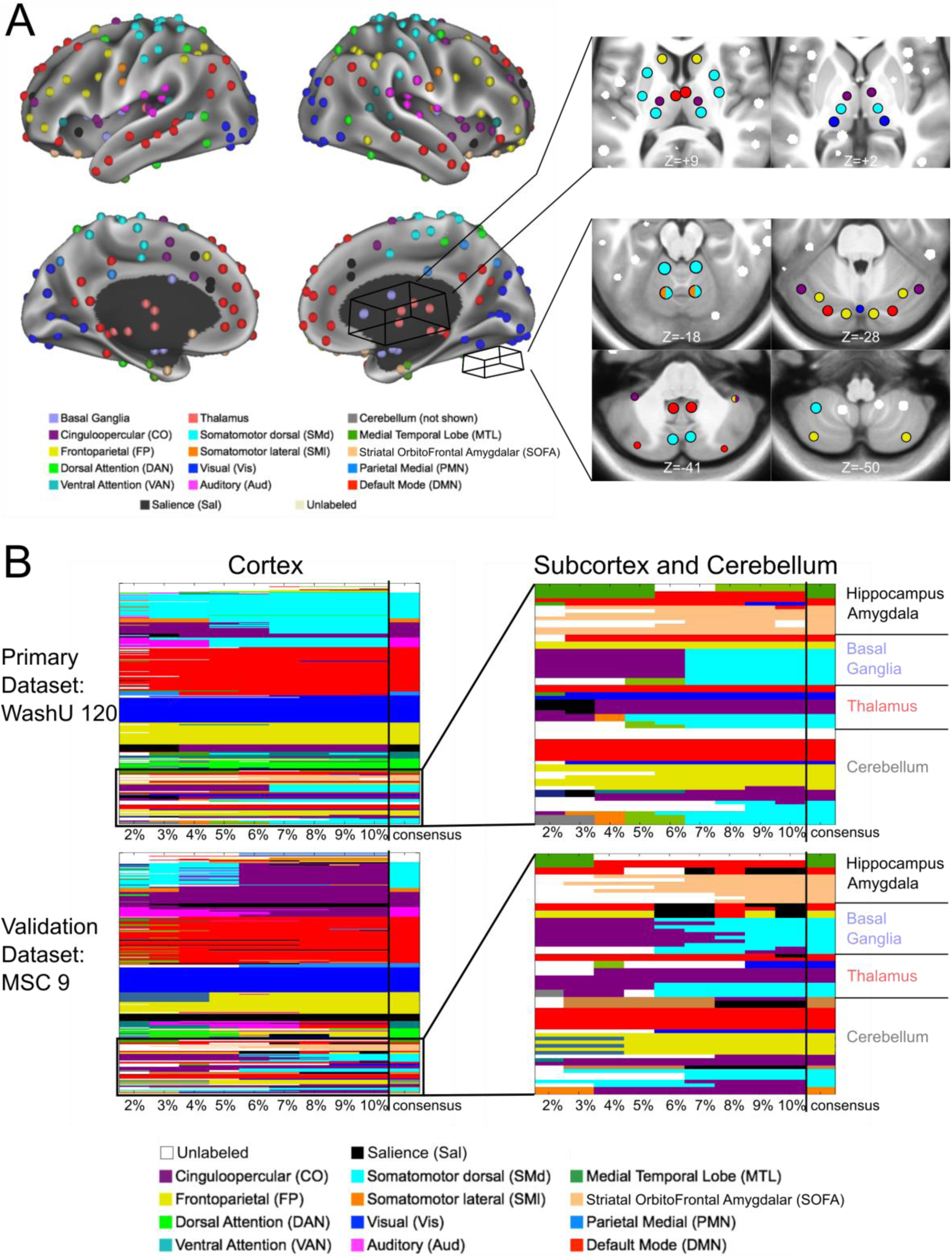
InfoMap-defined functional network communities. The InfoMap-defined functional network community of each ROI is displayed. (A) Cortical ROIs are shown projected onto the surface of the brain, and some of the non-cortical ROIs are displayed in axial slices to the right of the cortical surface. (B) The matrices represent the functional network assignment of each ROI across all tested edge densities (each column, denoted by the tick marks, represents one edge density), with the consensus functional network community displayed in the last column of each matrix (delineated by the vertical black line). Results are shown for the primary and validation datasets. The matrices on the left represent the cortical ROIs, and the colors correspond to the labels in A. The matrices on the right show zoomed-in results for all non-cortical ROIs. Results were highly consistent in the subcortex, cerebellum, amygdala, and hippocampus, with a total of 3 disagreements between datasets (in addition to 3 unlabeled ROIs at the bottom of the cerebellum forming their own “network” in the MSC dataset).

In the subcortex and cerebellum, the consensus network communities were as follows: ROIs in the caudate associated with the default mode network (head) or the frontoparietal network (tail). The putamen and globus pallidus ROIs joined the somatomotor dorsal network. In the thalamus, the default mode network was assigned to mediodorsal region, the cinguloopercular network to the laterodorsal and ventral anterior regions, the somatomotor dorsal network to the ventrolateral and ventral posterolateral regions, and the visual network to the ROI that includes the lateral geniculate nucleus and the posterior portion of the pulvinar. We use the names of the thalamic nuclei for convenience here, even though the ROIs encompass more gray matter than just the nuclei themselves. Cerebellar ROIs joined various networks, including the default mode, frontoparietal, and cinguloopercular networks (lateral), the somatomotor networks (motor cerebellar cortex), and the visual network (vermis). Most of the observed network assignments agree with known brain function, such as the association between ventral posteriolateral thalamic region and the somatomotor dorsal network.

While some ROIs did not vary in network membership across thresholds (e.g., the tail of the caudate ROIs), others changed network membership after a certain threshold (e.g., the putamen ROIs) or switched between two or more networks (e.g., some of the thalamic ROIs). This variation is similar to the variation seen with cortical ROI assignments (e.g., see Figure 1 from Power et al., 2011 and Figure 2A from Power et al., 2013) and is indicative of the loss of some finer-scale community structure at denser thresholds.

Importantly, we replicated these community assignments in the validation dataset (MSC; note that we did not use the secondary dataset for this analysis due to poor signal-to-noise in the subcortex). The consensus communities from the primary and validation datasets were broadly consistent across the two ROI Sets, with 55 out of 61 subcortical (including the amygdala and hippocampus) and cerebellar ROIs receiving the same assignment. For 24 ROIs, the InfoMap and winner-take-all assignments differed, requiring adjudication. A template-matching approach was implemented to adjudicate the consensus network of these ROIs. As an example, an ROI in the dorsal striatum was assigned to the salience (winner-take-all) and default mode (InfoMap) networks. The template-matching approach provided evidence for a final assignment to the salience network (Figure 7; see section 2.6.4; dice overlap with salience network template = 0.78).

**Figure 7:**
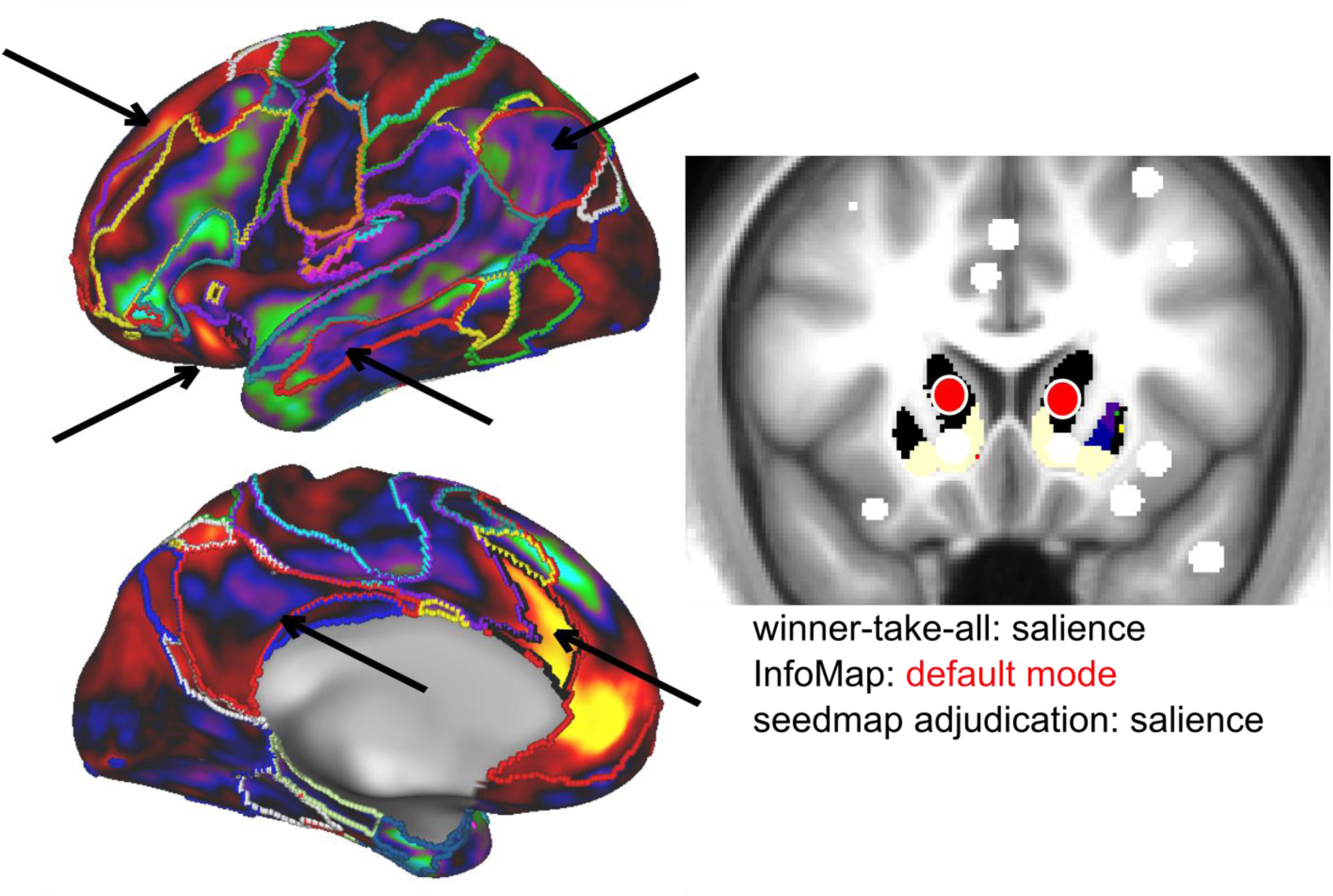
Disambiguation of discrepancies between assignments. In cases where the winner-take-all assignment and InfoMap solution differed, a template-matching approach (Gordon et al., 2017a) was used to determine the consensus ROI assignment. This exemplar ROI (head of the caudate) was assigned to the salience network (black) via the winner-take-all approach and the default mode network (red) via InfoMap. The ROI’s seedmap is more similar to the salience network (left, black outline) than the default mode network (left, red outline), especially on the lateral surface of the brain. Arrows highlight functional connectivity within the salience and default mode network regions. The template-matching approach confirmed the stronger similarity to the salience network (dice overlap with salience network template = 0.78).

Most cortical ROIs retained their functional network membership from Power et al., 2011 (ROI Set 1) or Gordon et al., 2016 (ROI Set 2). Nonetheless, with to the addition of the new ROIs, we observed two functional networks not previously observed with the original ROI sets: (1) a network composed of ROIs in the amygdala, ventral striatum, and orbitofrontal cortex, which we will call the “striatal-orbitofrontal-amygdalar (SOFA)” network and (2) a network composed of ROIs in the anterior hippocampus and entorhinal cortex, which we will call the medial temporal lobe (MTL) network. In addition, in ROI Set 1, 10 previously unlabeled ROIs were now assigned to a network: 4 to the SOFA network, 3 to the MTL network, 2 to the visual network, and 1 to the dorsal attention network. Also, 12 ROIs changed network membership: 2 from the cinguloopercular network to the somatomotor dorsal network, 1 from the auditory network to cinguloopercular network, and 9 from the salience network to the frontoparietal (2), dorsal attention (1), and cinguloopercular (6) networks. For ROI Set 2, 39 previously unlabeled ROIs were assigned to a network: 8 to the SOFA network, 10 to the MTL network, 16 to the parietooccipital network, and 5 to the default mode network (SI Figure 5). Again, consensus communities from the primary and validation datasets were broadly consistent.

Given the evidence that some regions in the subcortex connect to multiple functional networks (Greene et al., under review; Garrett et al., 2018; Hwang et al., 2017), we identified ROIs that overlap with integrative subcortical voxels (i.e., voxels with strong functional connectivity to more than one cortical functional network). We found that 8 ROIs in the basal ganglia and 6 ROIs in the thalamus contained a majority of integrative voxels (Figure 8). Therefore, we flag these ROIs as “integrative” and report the percent of integrative voxels in addition to their community assignment in the publicly available ROI list. In the cerebellum, no ROIs met these criteria.

**Figure 8:**
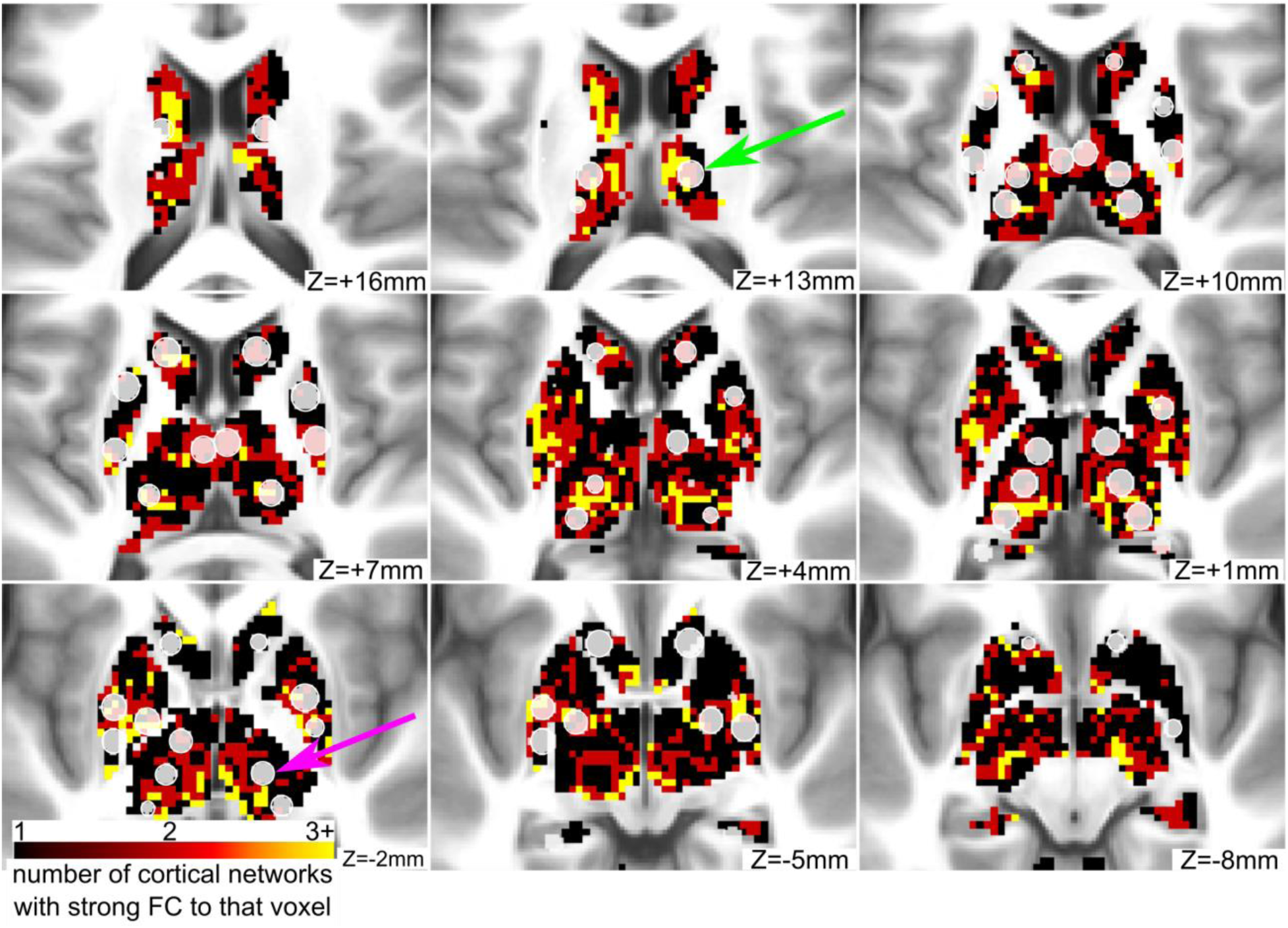
Some ROIs in the basal ganglia and thalamus have strong connectivity to multiple networks. Integrative voxels in the basal ganglia and thalamus are displayed in red and yellow colors. These voxels have strong connectivity to more than one cortical functional network. Some of the novel ROIs (displayed in translucent white) contain a majority of integrative voxels (green arrow), while other ROIs contain few or none (purple arrow).

### 3.4. Subcortical and cerebellar ROIs affiliate with known functional networks

To visualize the ROIs in functional network space, we created spring-embedded graphs, displayed in Figure 9 (other edge densities in SI Figure 6). The implemented spring model aggregates nodes with strong correlations between themselves and weak correlations with other nodes. Thus, it is possible to observe which nodes segregate into separate communities and which nodes act as connector hubs, mediating interactions across different network communities (Cohen and D’Esposito, 2016; Gordon et al., 2018; Gratton et al., 2012; Hagmann et al., 2008; Mattar et al., 2015; Power et al., 2013; van den Heuvel and Sporns, 2013).

**Figure 9:**
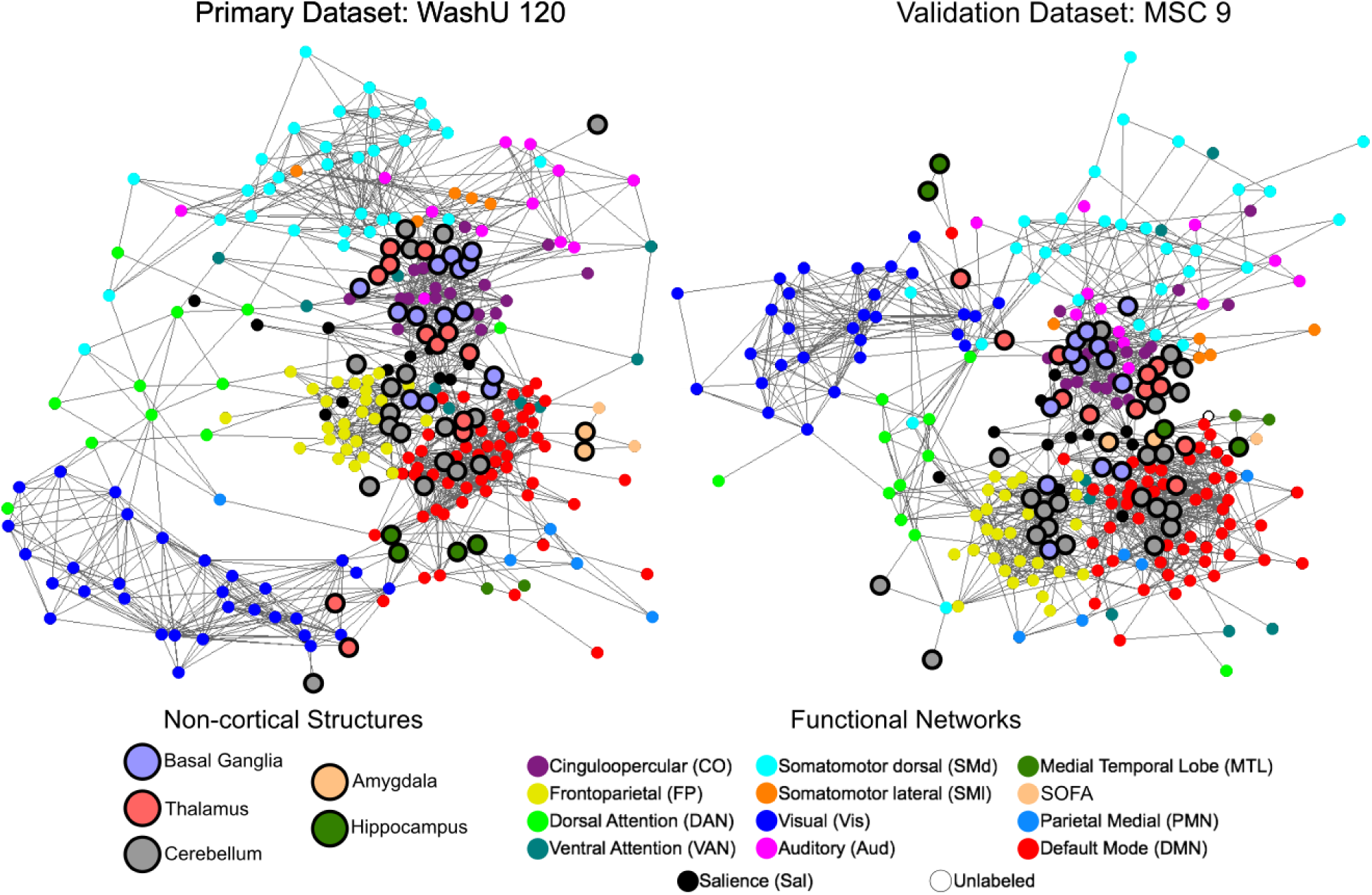
Spring-embedded graphs show that subcortical and cerebellar ROIs affiliate with well-characterized network communities. Spring-embedded graphs are displayed for ROI Set 1 using the primary and validation datasets at a structure-specific edge density threshold of 3% (other edge densities shown in SI Figure 6; see Section 2.6.3 for the thresholding procedure). Non-cortical ROIs are larger and have a bold outline. The color of each ROI represents its consensus functional network community assignment, except for the non-cortical ROIs, which are labeled by anatomical structure. The basal ganglia, thalamus, and cerebellum distribute throughout the graph, affiliating with well-characterized networks rather than segregating into their own communities.

As is evident from the position of the bolded network nodes, the subcortical and cerebellar ROIs were distributed throughout the spring-embedded graph. For instance, the cerebellar ROIs (gray) were not segregated from the rest of the network communities as in previous reports (Gratton et al., 2018a; Power et al., 2011). This finding was consistent between the primary and validation datasets. However, we observed that the basal ganglia, thalamus, and cerebellum did segregate into their own network communities when the graph was created without structure-specific edge density thresholding (SI Figure 6; see Section 2.6.3 for the thresholding procedure). That is, the basal ganglia, thalamus, and cerebellum clustered into their own separate network communities with standard edge density thresholding (applying the threshold uniformly to the whole correlation matrix), likely because of the lower correlation magnitudes associated with these regions.

To assess the effect of including the new ROIs on network topology, we examined two graph-theoretic network measures: modularity and participation coefficient (SI Figure 7; Rubinov and Sporns, 2010). Addition of the non-cortical ROIs decreased modularity, with structure-specific thresholding resulting in a further decrease. Similarly, the participation coefficient of ROIs in the subcortex was significantly higher, on average, than ROIs in the other structures, consistent with the finding that some of these ROIs contained integrative voxels. Structure-specific thresholding resulted in higher average participation coefficient for all structures.

## 4. Discussion

Here we present a set of regions of interest (ROIs) that sample the basal ganglia, thalamus, cerebellum, amygdala, and hippocampus more completely than previous ROI sets in order to provide a whole-brain description of functional network organization. We found that the refined region sets recapitulate previous network organization results in the cortex and extend functional brain network characterization to the subcortex and cerebellum. Notably, these results replicated across independent datasets. In addition, with the inclusion of the new ROIs, we observe two additional functional networks that were not present in Power et al. (2011) and Gordon et al. (2016): a striatal-orbitofrontal-amygdalar (SOFA) network and a medial temporal lobe (MTL) network.

### 4.1. Improved sampling of the subcortex and cerebellum

Many recent research efforts have used the 264 ROIs from Power et al., 2011 or the 333 surface-based parcels from Gordon et al., 2016 to study brain network organization. These studies have examined both structural and functional network organization in a wide variety of samples, including healthy young adults (Power et al., 2013; Zanto and Gazzaley, 2013), developmental cohorts (Gu et al., 2015; Nielsen et al., 2018; Rudolph et al., 2017), older adults (Baniqued et al., 2018; Gallen et al., 2016), and a plethora of neurological and psychiatric populations (Gratton et al., 2018a; Greene et al., 2016; Sheffield et al., 2015; Siegel et al., 2018). We have gained a better understanding of typical and atypical human brain organization from these efforts. However, a full characterization of whole-brain network organization in these populations is incomplete due to the common unsderrepresentation of the subcortex and cerebellum. While there is recent work that has focused separately on networks in the thalamus, subcortex, and cerebellum (e.g., Bell and Shine, 2016; Buckner et al., 2011; Choi et al., 2012; Greene et al., 2014; Hwang et al., 2017), here we offer a set of ROIs that encompass all of these structures to encourage broader adoption of a whole-brain approach (as recently used in (Nielsen et al., 2019)).

The functionally-defined subcortical and cerebellar ROIs presented in the current work provide a better sampling of these structures. By improving their representation, we were able to delineate well-characterized and additional functional network communities (relative to our past descriptions). The ability to uncover these networks, which have been previously described using other methods, illustrates the importance of representing the entire brain in network-based analyses. Further, these improved ROI sets may allow future studies to discover previously unobserved, yet critical deviations in functional network organization in diseases and disorders in which the subcortex and cerebellum are implicated (e.g., Parkinson Disease, Tourette Syndrome, Schizophrenia).

It is worth noting that, by definition, the cortical surface parcels omit the subcortex and cerebellum. Yet, it is technically possible to parcellate the subcortex and cerebellum using an adapted gradient-based methodology (such as the one from Gordon et al., 2016). This approach would require extending the gradient technique to three dimensions. As fMRI technology and analysis strategies improve, it would be useful to compare the current results to a full subcortical and cerebellar parcellation using this or other gradient-based techniques.

A methodological issue to note is that the winner-take-all approach used to define subcortical and cerebellar parcels here may be sensitive to the number of *a priori* cortical networks used in the analysis. Increasing the number of cortical networks included may allow for finer parcellation of certain structures. Here we used previously well-characterized cortical networks that have been consistently found using multiple methods by multiple research groups (Power et al., 2011; Yeo et al., 2011). Conversely, InfoMap does not require an *a priori* number of networks, but may be sensitive to thresholding issues (discussed below). Importantly, final network assignments for the ROIs were designated using a combination of both techniques. Moreover, 14 ROIs that demonstrated strong connectivity with multiple functional networks were noted as integrative.

### 4.2. Functional connectivity of the refined ROIs is consistent with previous studies and replicates across independent datasets

Correlation seedmaps from the refined ROIs agree with functional connectivity profiles reported in previous studies. For example, the ROIs added to the ventral striatum and the head of the caudate correspond closely to the seeds placed in the superior ventral striatum (VSs) and dorsal caudate (DC) reported in Di Martino et al., 2008, and our seedmaps are highly similar to theirs. Likewise, seedmaps from the amygdala agree well with those from Roy et al., 2009. The same is true for the thalamus (Hwang et al., 2017) and cerebellum (Buckner et al., 2011).

Moreover, the full correlation structure (shown in correlation matrices) was quite comparable across the diverse datasets. The one major discrepancy was that in the subcortical portion of the matrix from the secondary (HCP) dataset, we observed correlations near zero. The reason for this observation is likely poor temporal signal-to-noise ratio (tSNR) in the subcortex of HCP data (Ji et al., 2019). Several factors may contribute to this poor tSNR. (1) The HCP used a custom scanner and coil, which caused unique magnetic field inhomogeneities, possibly in part due to subjects’ heads being outside of the isocenter of the field. (2) The imaging sequence used an aggressive multiband factor and TR (MB = 8, TR = 0.72s) and (3) small voxels (2 cubic mm) were used for acquisition (Glasser et al., 2013; Van Essen et al., 2012). Each of these factors substantially increase electronic, thermal, and other physical sources of noise (Triantafyllou et al., 2005) relative to slower sequences with larger voxels. These effects may be amplified as a function of the distance of the imaged structure from the head coil, resulting in the poorest tSNR in the subcortex. Further work is needed to determine the specific contributions of each factor, as well as others heretofore unconsidered, to the observed poor tSNR.

The presented group-level descriptions converge on a very similar picture of functional network organization in the subcortex and cerebellum. However, there are individual differences in both subcortical and cerebellar functional network organization (Marek et al., 2018; Greene et al., under review), as have been found in cortical functional network organization. Future work designed for in-depth study of individuals, as in Poldrack et al., (2015), Filevich et al., 2017, Braga and Buckner (2017), and Gordon et al. (2017b), will be important for elucidating such individual differences. In fact, in-depth study of the cerebellum (Marek et al., 2018) and subcortex (Greene et al., under review) in individuals reveals both common and unique features in its functional organization. Furthermore, future work may be able to include the brainstem as well in a whole-brain functional network atlas, although there are several technical issues to overcome (e.g., CSF pulsations, small nuclei size).

### 4.3. SOFA and MTL functional networks map onto known human brain systems

Group-average functional network organization in the cerebral cortex is largely consistent across studies (Power et al., 2011; Yeo et al., 2011), and the addition of refined subcortical and cerebellar ROIs did not change functional network organization in the cortex substantially (although we observed associations between these canonical networks and ROIs in the subcortex and cerebellum). However, the addition of these subcortical and cerebellar ROIs allowed for the identification of two additional functional networks compared to the networks reported using the original ROI sets in Power et al. (2011) and Gordon et al. (2016): (1) the “SOFA” network composed of the amygdala, orbitofrontal cortex, and ventral striatum, and (2) the “medial temporal lobe (MTL)” network composed of the anterior hippocampus and entorhinal cortex. It is worth noting that the SOFA network has been observed in studies focusing on reward and emotion processing (Camara et al., 2009) and its cortical and striatal portions are very similar to the limbic network from Yeo et al., (2011) and Choi et al., (2012). The MTL network has been observed in a study of highly-sampled individuals (Gordon et al., 2017b) as well as studies focused on the hippocampus (Greicius et al., 2009). Here, we demonstrate that these networks are measurable at the group-level when the whole brain is represented sufficiently. In addition, we found that some cortical ROIs that were previously unlabeled (i.e., they did not group with any community) received labels with the inclusion of the refined subcortical and cerebellar ROIs, with many of them joining the SOFA and MTL networks.

The SOFA and MTL functional networks map onto well-characterized brain systems. Most of the ROIs in the SOFA network are likely connected to each other anatomically in rodents, nonhuman primates, and humans (Ongur and Price, 2000; Carmichael and Price, 1995; Amaral and Price, 1984). Moreover, these brain areas are known to be functionally related, as they are important for various aspects of decision making and reward-related behavior, such as economic choice (Padoa-Schioppa and Assad, 2006), emotional regulation (Phelps, 2006), and gambling (Bechara et al., 2000, 1997). Likewise, the ROIs in the MTL network are well-connected anatomically (Duvernoy, 1988; Woolsey et al., 2008) and support various aspects of memory formation, consolidation, and retrieval, as well as other important functions, such as spatial mapping (Burgess et al., 2002; Moser and Moser, 1998; Tulving and Markowitsch, 1998). Though our current work is agnostic to the function of these brain systems, we show that their constituent regions demonstrate coherent spontaneous fluctuations in infraslow BOLD signal.

### 4.4. Some ROIs in the subcortex exhibit functional connectivity with multiple cortical networks

There is evidence to suggest that regions of the basal ganglia and thalamus act as integration zones, combining information from multiple functional brain networks (Garrett et al., 2018; Haber, 2003; Hwang et al., 2017). A majority of the non-cortical ROIs included in this study exhibited functional connectivity primarily with just one cortical functional network, and therefore, can be considered network-specific ROIs. Particularly in the cerebellum, we found that all ROIs were network-specific. However, in the basal ganglia and thalamus, some of the ROIs overlapped with integration zones, as they contained voxels with strong connectivity to multiple cortical networks. These subcortical integration zones have been explored more fully in Greene et al., (under review), demonstrating preferential integration of particular networks with given subregions of the basal ganglia and thalamus. These findings are consistent with models of subcortico-cortical connectivity describing parallel and integrative circuits (Haber, 2003) as well as with previous neuroimaging work showing that subcortical structures contain integrative hubs (Garrett et al., 2018; Hwang et al., 2017).

We allow users of our ROI Sets to account for the above feature of brain organization as flexibly as possible. For each ROI, we provide the percent of voxels overlapping with integrative zones, and we flag “integrative” ROIs as those with a majority of voxels in such regions (in our publicly available lists). Thus, users can incorporate this information as desired and include or exclude certain ROIs in order to craft an ROI Set best-suited to investigate their research questions.

### 4.5. Subcortical and cerebellar ROIs affiliate with known functional networks

To visualize the organization of the ROIs in functional network space, we created spring-embedded graphs. We observed that the subcortical and cerebellar ROIs affiliate with various well-characterized network communities composed of cortical regions instead of segregating on their own (i.e., away from cortical ROIs), particularly after structure-specific thresholding (see below). This organization fits with the known anatomy and function of the subcortex and cerebellum better than a model in which each structure is segregated into its own community. For instance, individual nuclei in the thalamus project directly to distinct brain systems (Woolsey et al., 2008) and play unique roles in behaviors associated with those systems (Guillery, 1995; Van Der Werf et al., 2000). Likewise, striato-cortical and cerebello-cortical anatomical connections show specific projections to unique regions of cortex (Woolsey et al., 2008) and are known to be integral for the function of various large-scale, distributed systems, such as the motor system (Glickstein and Doron, 2008) and regions of higher order systems (Alexander et al., 1986; Strick et al., 2009). Investigation of network measures revealed that the subcortical ROIs have a higher participation coefficient, on average, than other structures, meaning they have modest-to-high correlations with multiple networks. This result is consistent with the idea that subcortical structures contain integrative hubs (Hwang et al., 2017); Greene et al., under review). Likewise, the non-cortical ROIs decrease the modularity of the whole network, reflecting decreased segregation and increased integration. These results are consistent with the finding of integrative ROIs in the subcortex described above. It should be noted, however, that several methodological factors may affect or potentially bias network measures like participation coefficient, such as structure-specific thresholding and the interaction between structure size and BOLD fMRI spatial autocorrelation.

The demonstration that subcortical and cerebellar ROIs affiliate with cortical networks was revealed by the use of structure-specific edge density thresholding (i.e., thresholding the cortex, subcortex, and cerebellum separately). In most network analyses, only the strongest positive correlations are considered for network-based analyses, such as spring-embedded graphs. However, subcortical correlations are generally weaker than cortical correlations (likely due to distance from the head matrix coil and signal dropout due to sinuses). Thus, if the top 5% strongest positive correlations are selected, almost all subcortical correlations will be excluded. To avoid this exclusion, we implemented structure-specific thresholding. This choice ultimately affects the nature of the spring-embedded graph as well as the determination of functional network communities and network measures. Without structure-specific thresholding, subcortical ROIs group with one another into two separate network communities (basal ganglia and thalamus), while the entire cerebellum is lumped into one network community. In terms of human brain functional organization, this pattern of clustering seems artificially inflated due to low subcortex-to-cortex and cerebellum-to-cortex correlations. By using structure-specific thresholding, we were able to observe functional network organization that is more consistent with the known functions of the subcortex and cerebellum. However, this approach may affect graph theoretic network measures in the subcortex and cerebellum, and thus, deserves future investigation.

### 4.6. “Optimal” ROI set depends on research question

There are advantages to both anatomical and functional network-based divisions of ROIs. For instance, anatomical network divisions allow for analysis of important distinctions between the cortex, subcortex, and cerebellum, whereas functional network divisions are likely to better represent putative brain function. Likewise, there are fundamental differences between anatomical and functional atlases, with anatomical atlases parsing the brain according to anatomical divisions and cytoarchitecture, and functional atlases parsing the brain according to functional criteria. For instance, the cerebellum is probabilistically divided into lobes and crura in one anatomical atlas (Diedrichsen et al., 2009). While many of these divisions align well with the ROIs presented here, some ROIs do not fit within the probabilistic boundaries. Thus, some ROIs may better reflect functional rather than anatomical divisions in the cerebellum. Similarly, many of the divisions in a commonly used subcortical anatomical atlas (Morel, 2013) agree well with the ROIs presented here, with the exception of a few anatomical parcels that are likely beyond the resolution of the fMRI data used here. Ultimately, researchers should be cognizant of these effects when choosing how to perform network-based analyses and which atlas or ROI Set to use. We advise readers to use the analysis strategy and atlas that best suits their research questions.

### 4.7. Conclusions

We created new subcortical and cerebellar ROIs to improve the representation of these structures for brain network analysis. Combining these new ROIs with previously characterized cortical ROIs allowed further insight into whole-brain functional network organization. Going forward, inclusion of these ROIs will yield more comprehensive results from fMRI studies of typical and atypical brain organization and function. The ROI Sets and consensus functional network assignments described here are available for immediate download and use at https://greenelab.wustl.edu/data_software.

## Acknowledgements

This work was supported by National Institutes of Health grants T32 NS073547 (BAS), F32 NS092290 (CG), T32 MH10001902 (SM), K23 NS088590 (NUFD), UL1 TR000448 (NUFD), R01 NS32979 (SEP), R01 NS06424 (SEP), and K01 MH104592 (DJG); the National Science Foundation grant DGE-1745038 (RVR); the Jacobs Foundation grant 2016121703 (NUFD); the Child Neurology Foundation (NUFD); the Mallinckrodt Institute of Radiology grant 14-011 (NUFD); the Hope Center for Neurological Disorders (NUFD, BLS, SEP); the James S. McDonnell Foundation Collaborative Activity Award (SEP); the McDonnell Center for Systems Neuroscience (NUFD, BLS, DJG); and, the Tourette Association of America (DJG).

## Author Contributions

BAS, CG, and DJG designed the study. BAS, CG, SM, RVR, and DJG analyzed the data. All authors wrote the manuscript.

## Supplemental Figures

**SI Figure 1:**
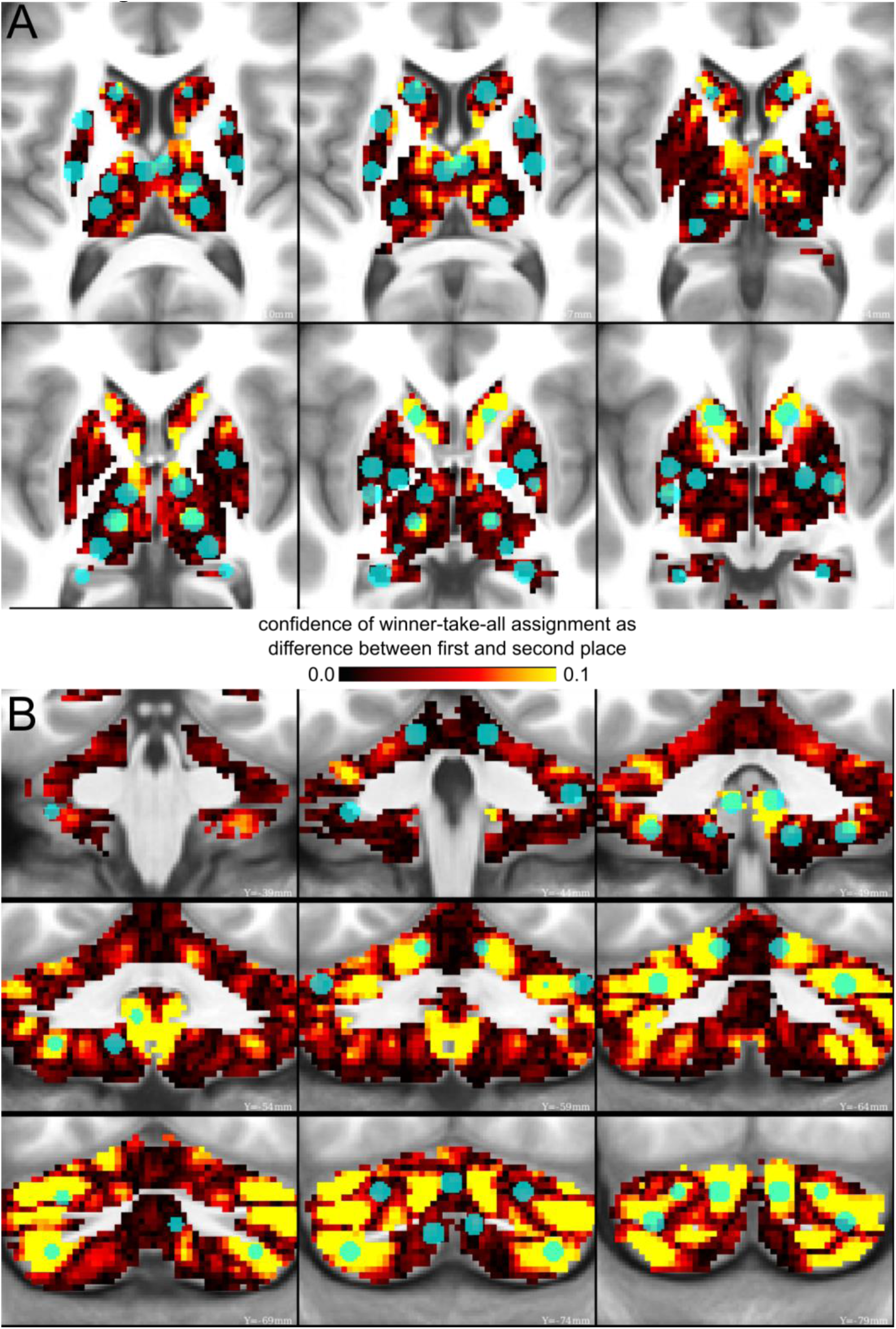
ROIs in high confidence winner-take-all parcels. The difference between the top two functional networks from the winner-take-all parcellation is displayed. Warmer colors indicate high confidence winner-take-all network assignments. The ROIs are overlaid in a translucent blue. We observed that 35 out of 55 (64%) ROIs in the basal ganglia and thalamus (A) and cerebellum (B) contained “clear winner” voxels (i.e., voxels with a difference in correlation strength between the first and second place networks ≥0.05). The remaining 20 ROIs contained voxels with strong functional connectivity to multiple networks, suggesting that they may act as integrative hubs (Hwang et al., 2017). An alternative interpretation is that the assignments for these 20 ROIs are low confidence (Marek et al., 2018). To allow for user flexibility, we created flags for these ROIs in the publicly available files, so users will be able to exclude them if desired.

**SI Figure 2:**
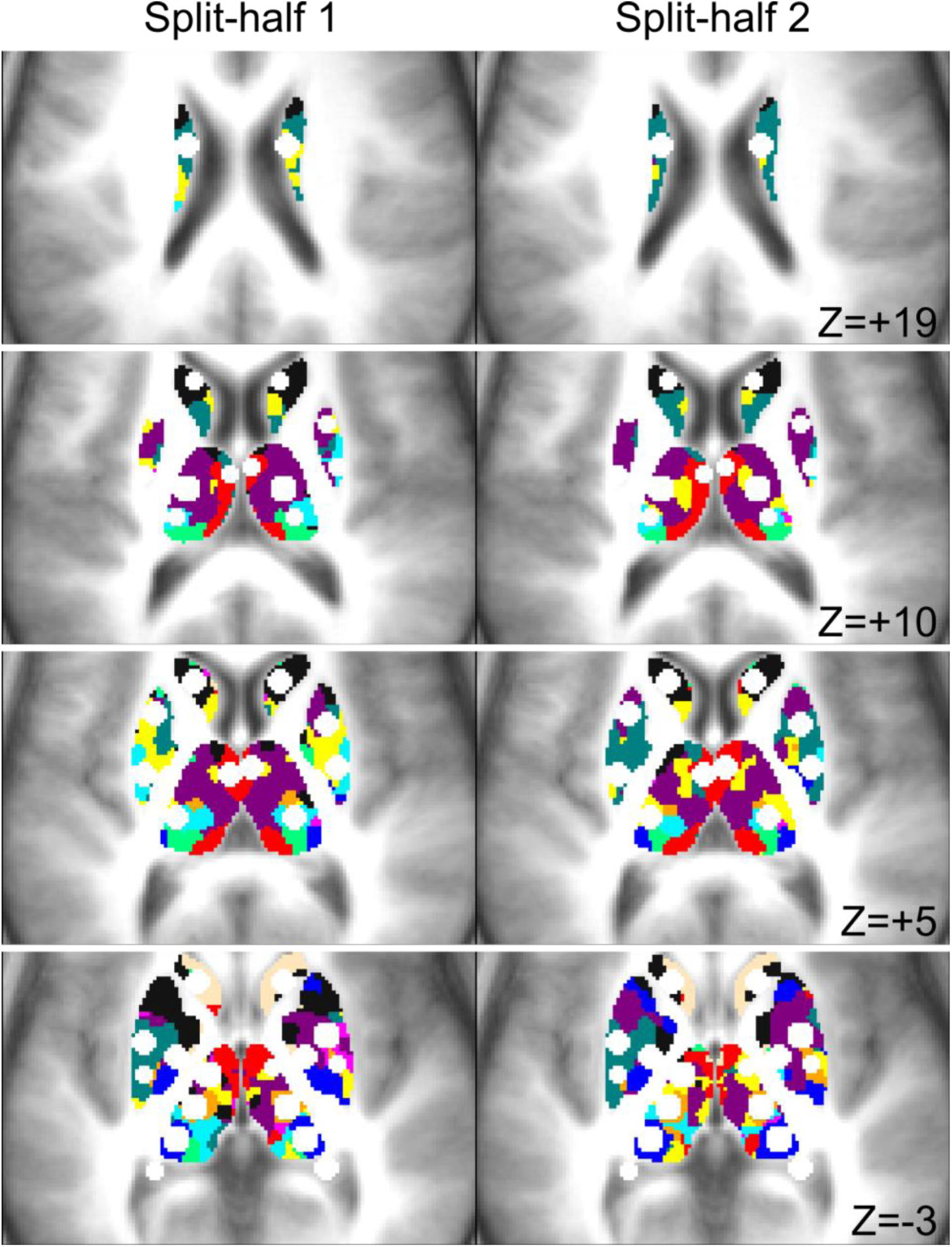
Consistency of winner-take-all assignment between split-halves. In locations where the ROIs are located, there is good consistency of the winner-take-all assignments in the two split-halves of the primary dataset (N=60 each).

**SI Figure 3:**
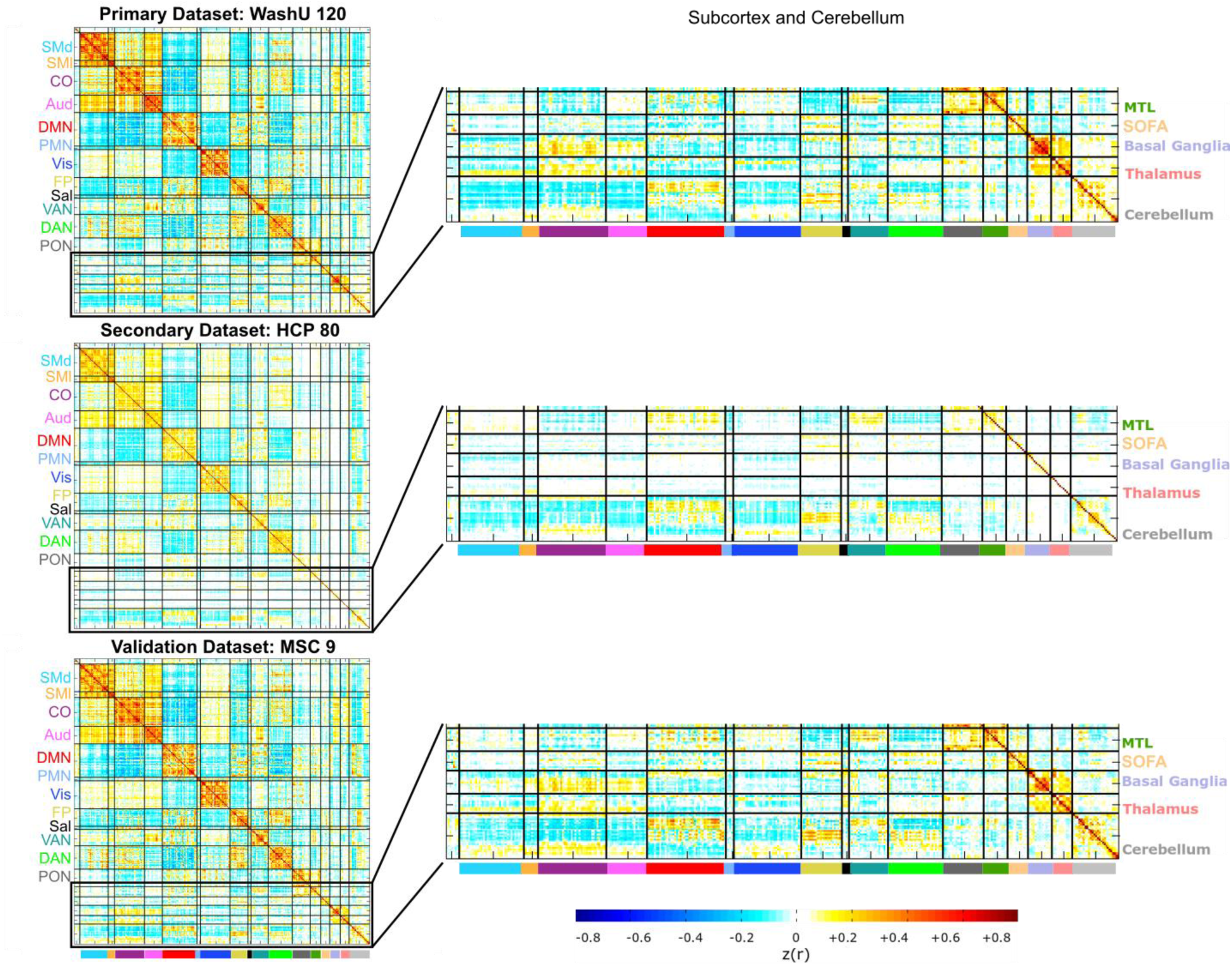
Correlation matrices for ROI Set 2. The center of each surface-based parcel from Gordon and Laumann et al., 2016 (*Cerebral Cortex*) was projected into volume space and combined with the new ROIs presented in this work to create ROI Set 2. The mean BOLD timeseries from all voxels within each ROI was extracted. The correlation matrices for each dataset are displayed, and zoomed-in portions of the matrix corresponding to the new ROIs are on the right.

**SI Figure 4:**
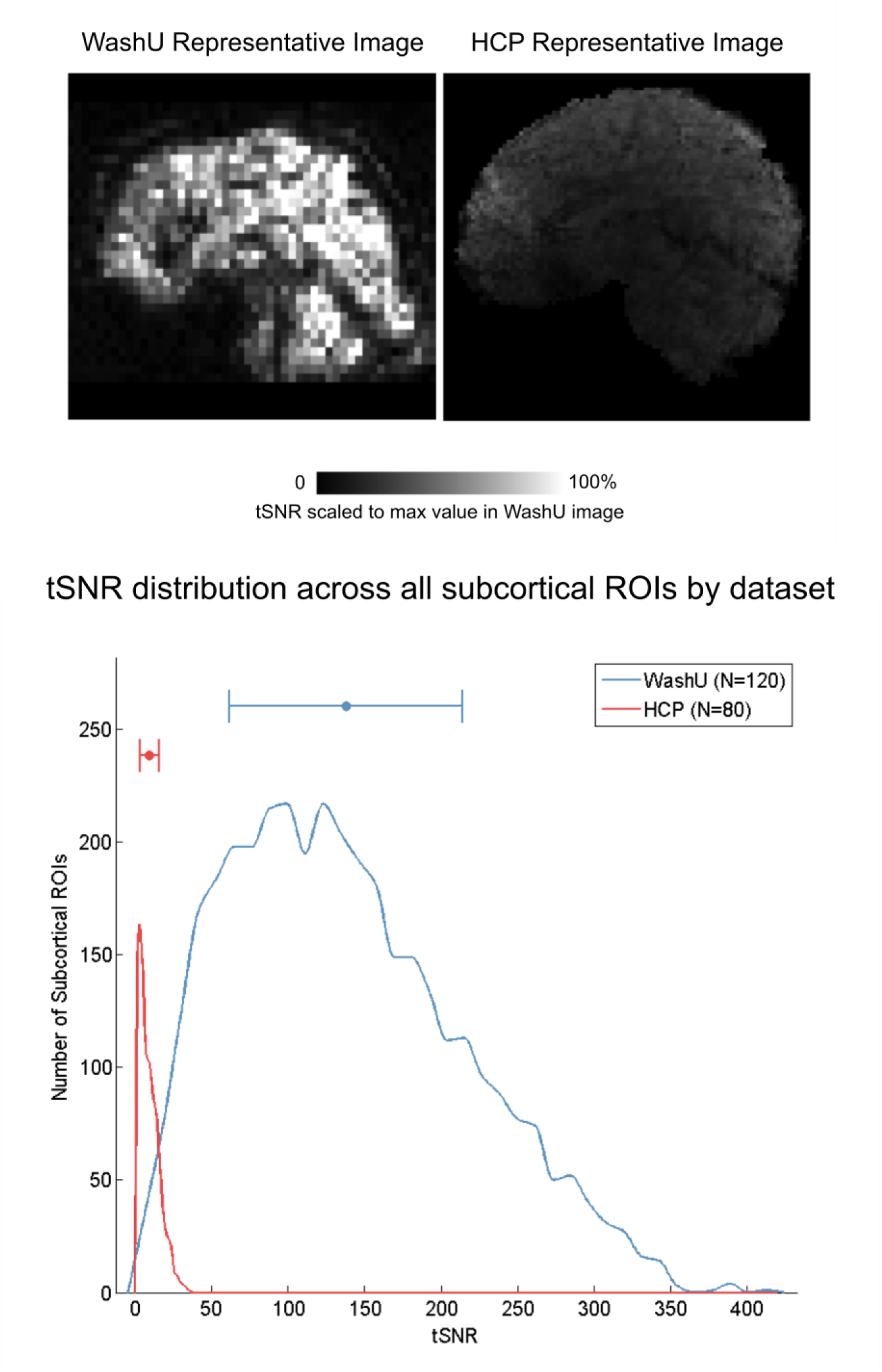
Poor temporal Signal-to-Noise Ration (tSNR) in the subcortex of Human Connectome Project data. Similar to previously published studies, we found that there was poor tSNR in the subcortex of HCP data. Representative images of tSNR (mean divided by standard deviation of the BOLD timeseries at each voxel) are displayed for an individual from the primary dataset (WashU 120) and from the HCP dataset (top). The images are scaled to the maximum tSNR value in the WashU image. The distributions on the bottom represent tSNR for all subcortical ROIs across each individual in each dataset. The distribution for HCP (red) is significantly worse than the primary dataset (mean +/- std = 9.63 +/- 6.48 for HCP; 137.79 +/- 76.22 for WashU).

**SI Figure 5:**
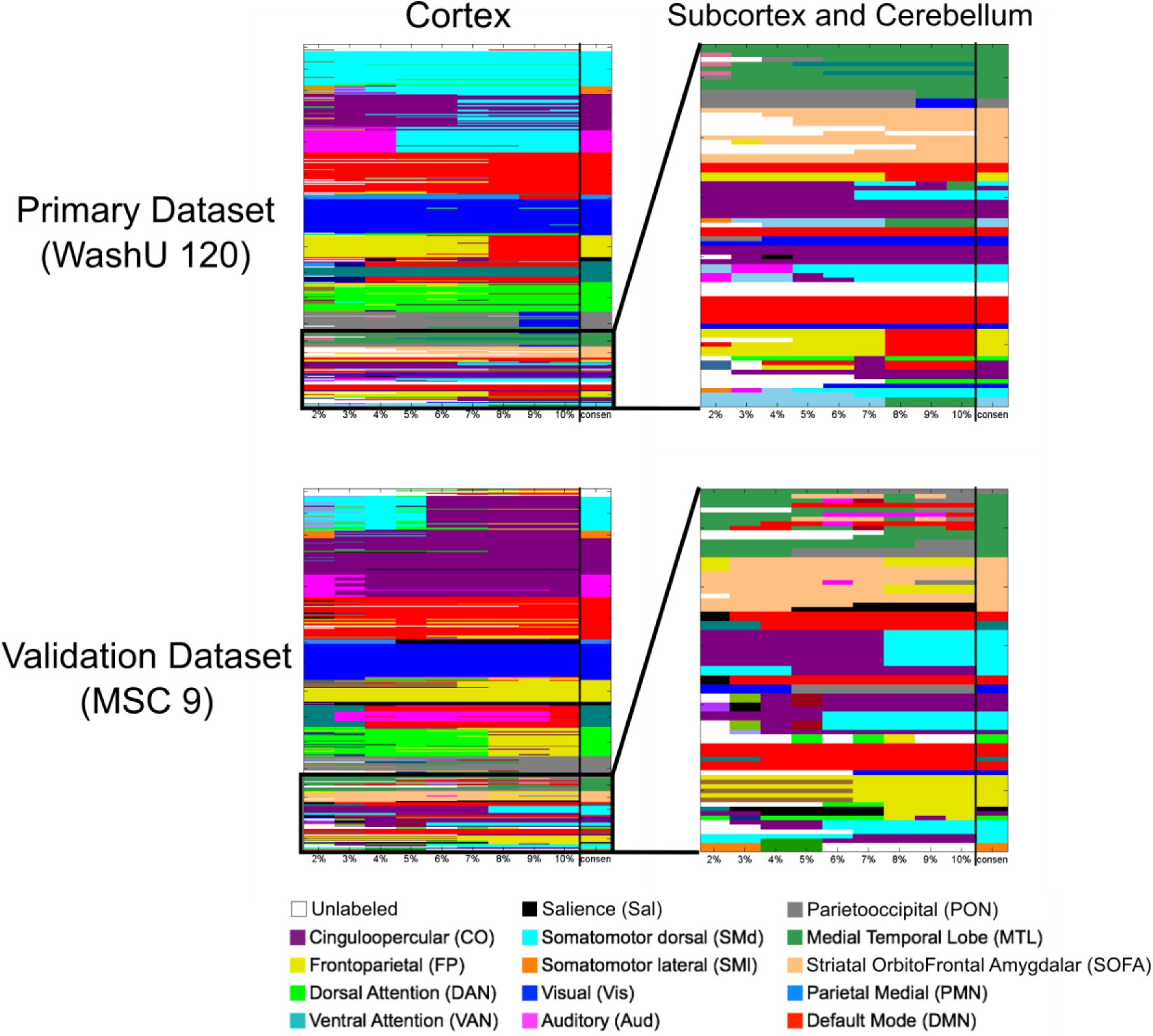
InfoMap-defined functional network community assignments for ROI Set 2. The functional network communities detected via InfoMap are displayed for the primary (WashU 120; top row) and validation (MSC; bottom row) datasets. The results were very similar to those shown in the main text, and there was good agreement between the two datasets. The primary difference is the presence of the Parietal Occipital Network in the cortex (gray), which was not observed with ROI Set 1.

**SI Figure 6:**
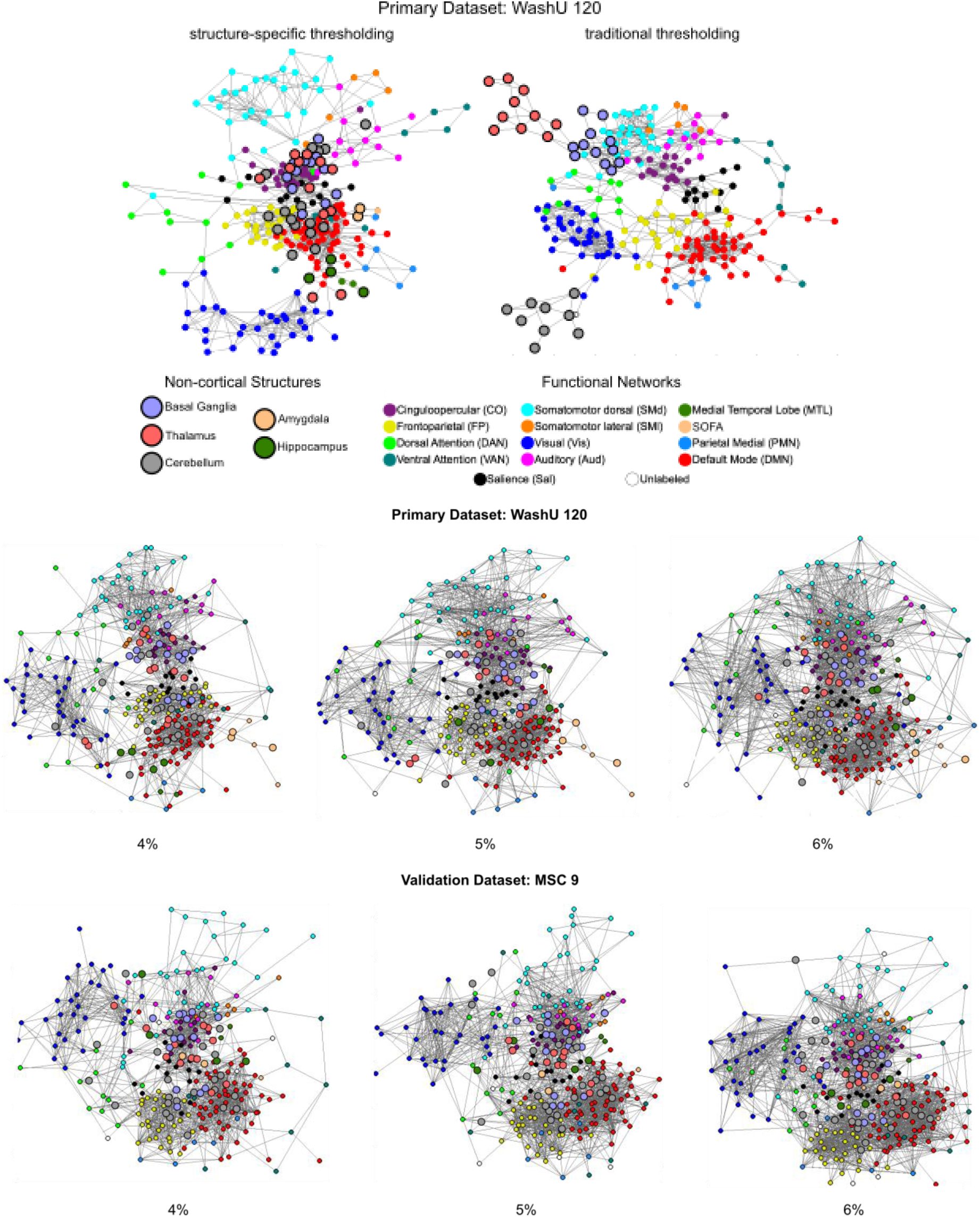
Spring-embedded graphs at other tested edge densities. The top portion of the figure shows the difference between structure-specific edge density thresholding and traditional thresholding (uniform across the matrix). The basal ganglia, thalamus, and cerebellum segregate into their own network communities when traditional thresholding is used (top right graph). Spring-embedded graphs for other structure-specific edge density thresholds are displayed below for the primary and validation datasets. The non-cortical ROIs (larger, bold outlines) distribute throughout each graph, integrating with known functional networks.

**SI Figure 7:**
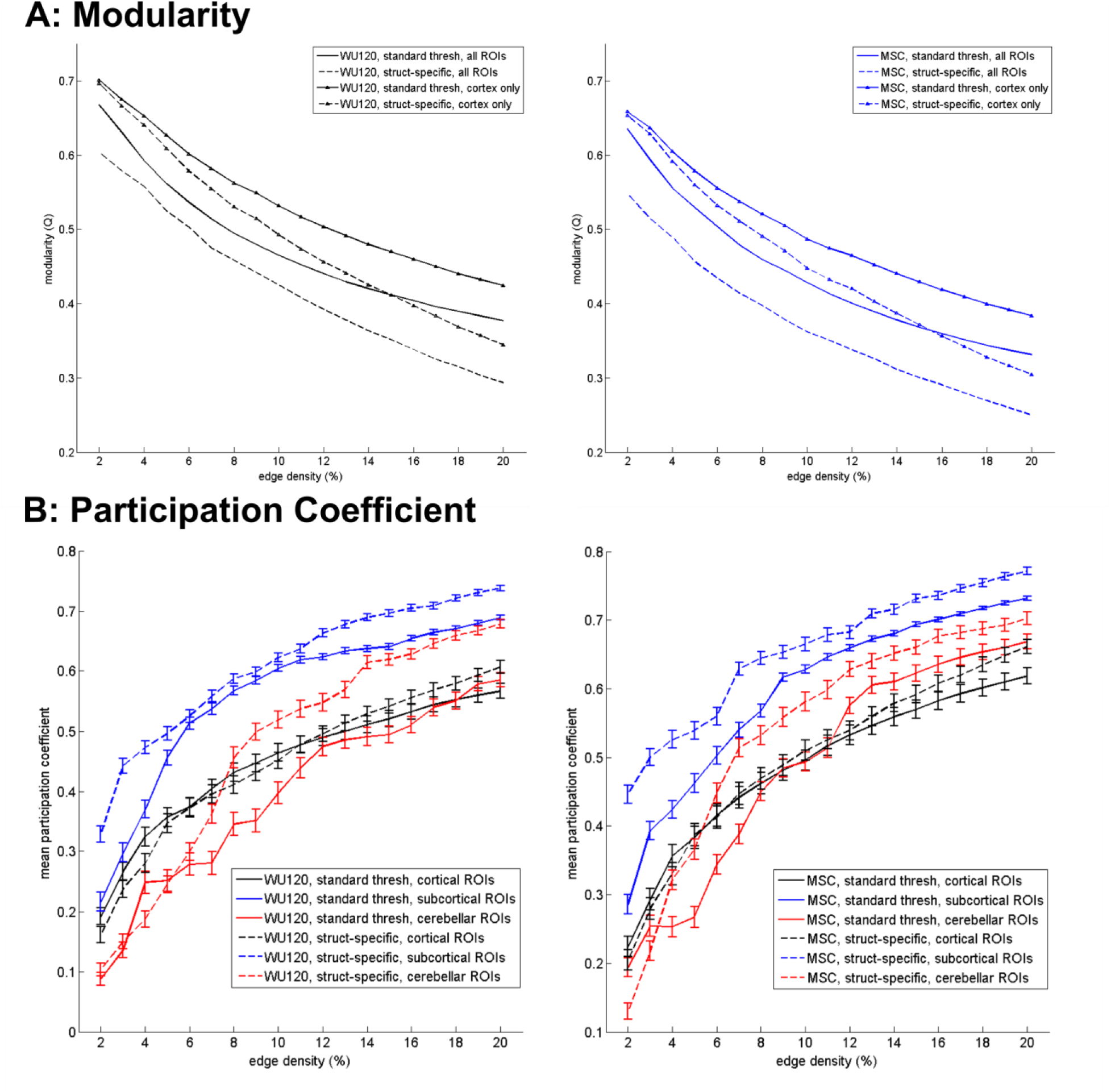
Graph-theoretic network measures. (A) Inclusion of non-cortical ROIs decreased modularity. Graphs display the modularity statistic for ROI Set 1 with (no marker) and without (triangle marker) the subcortical and cerebellar ROIs, as well as with (dashed lines) and without (full lines) structure-specific edge density thresholding, for the WashU 120 (black; left) and MSC (blue; right) datasets. Modularity was calculated always assuming the consensus network assignment across a variety of edge density thresholds. (B) Subcortical ROIs have higher average participation coefficient than cortical and cerebellar ROIs. Graph displays the average participation coefficient for all ROIs within a structure with (dashed lines) and without (full lines) structure-specific edge density thresholding. Participation coefficient was computed for each ROI and averaged across all ROIs in the cortex (black), subcortex (blue), and cerebellum (red) for the WashU 120 (left) and MSC (right) datasets. The result of this analysis is shown for a variety of edge densities while always assuming the consensus network assignments.

